# Multimodal temporal modeling of behavioral reorganization across threat learning and extinction in rats

**DOI:** 10.64898/2026.07.17.739137

**Authors:** Bin Yin, Jiaojiao Rao, X.T. (XiaoTian) Wang

**Affiliations:** School of Psychology, Fujian Normal University, Fuzhou, China; School of Humanities and Social Science, The Chinese University of Hong Kong (Shenzhen), Shenzhen, China

**Keywords:** fear conditioning, extinction, instrumental behavior, ultrasonic vocalization, hidden Markov model, functional recovery

## Abstract

After threat disrupts a learned activity, does engagement return as defensive responding later declines? Tri-reference-point theory describes decisions under risk relative to a minimum requirement, the status quo and a goal; we extended this logic to defense and engagement. We followed freezing, 22-kHz and 50-kHz calls, and food-reinforced lever responses across conditioning, extinction, delayed retesting and context changes in 29 male rats (45,300 five-second windows). A hidden Markov model described recurring response combinations, or configurations, and their persistence. For every animal, it predicted behavioral sequences excluded from fitting better than a matched model that ignored temporal order. Among models with two to eight configurations, eight predicted best; four provided a coarser description. This coarser model separated relatively brief press-active patterns from more persistent vocal-active and low-output patterns. Low-output patterns contained few calls and lever responses, but could involve either little or substantial freezing. In the repeated-session comparison, the estimated share of time in press-active patterns increased on average in the full extinction group (n=9) and decreased in cue-free context controls (n=8), from different first-session levels. These descriptive trajectories included renewed instrumental engagement alongside continued defensive immobility. We separately tested whether configuration summaries added predictive value beyond the measured behaviors for later responding or whole-recording fecal-pellet counts, finding no consistent benefit. Tracking how behavioral combinations change and persist provides a computational framework for evaluating functional adaptation beyond defensive decline.

**Impact statement:** Functional recovery after threat may reorganize an individual’s repertoire rather than reproduce it. Following instrumental engagement together with defense and vocal activity makes this distinction measurable, while temporal models describe their coordination.

**Plain language summary:** A rat can freeze less after danger yet still fail to resume an activity it has learned. Understanding recovery therefore means following how the animal uses opportunities to act during and after threat. We followed rats that had learned to press a lever for food through threat learning, repeated presentations of the threat cue without shock (extinction), and later tests in familiar or changed circumstances.

Comparison groups received no shocks or encountered the same chambers without the extinction sounds. During these stages, we recorded freezing, physical lever responses and two types of ultrasonic calls. A computer model linked combinations of these responses across successive five-second periods, asking how long each combination lasted and which followed it. Keeping this temporal order improved prediction of behavior compared with treating each period independently. Pressing formed relatively brief bouts, whereas patterns dominated by calling or low vocal and instrumental output persisted longer. The low-output periods were especially informative: some involved little freezing, while others involved substantial freezing. Across repeated sessions, pressing-related patterns increased on average in the extinction group and decreased in the cue-free comparison group, which started at a much higher level. High-freezing periods remained present. Renewed activity and continued defense could therefore be parts of the same changing repertoire, rather than opposite ends of a single fear scale. Finer models predicted the sequences better, but their summaries did not consistently improve prediction of later behavior or the number of fecal pellets counted after a session. The approach makes re-engagement visible in relation to pauses, calling and defense, providing a way to evaluate functional change without assuming that recovery must recreate the pre-threat pattern.

## Introduction

Defensive responses have provided much of the experimental foundation for understanding how animals learn about danger (1–3). Recovery, however, concerns what happens to the rest of the animal’s behavior when those responses decline. An animal may become less defensive without resuming a previously learned action. Renewed engagement, when it occurs, may combine familiar responses differently rather than reproduce the pre-threat repertoire. Recent temporal ethograms show why this distinction matters: conditioned threat cues organize several defensive and orienting behaviors while suppressing reward seeking (4, 5). The functional consequences of threat therefore extend beyond the magnitude of a single fear response. Understanding those consequences requires following what the animal does instead, both while danger is present and after its immediate expression subsides. We refer to this organization of competing behavioral priorities as fear-goal arbitration, without assuming a particular decision mechanism.

The first requirement is to observe responses that can change together or dissociate. Freezing provides an established measure of conditioned defense, but it does not exhaust the ways threat can affect behavior. Periaqueductal gray lesions can disrupt conditioned freezing while sparing conditioned suppression of instrumental behavior (6); reduced reward seeking is therefore not simply freezing by another name. Ultrasonic vocalizations (USVs) add information that neither movement nor task performance supplies. Calls near 22 kHz occur in aversive and alarm contexts, whereas 50-kHz calls occur across social, rewarding, and novel situations (7–10). Their interpretation depends on context and acoustic subtype: pooled 50-kHz output cannot by itself identify exploration or positive affect. Food-reinforced lever pressing measures engagement in a learned instrumental task. Recording these channels together makes it possible to distinguish changes in defense, vocal activity, and instrumental responding without requiring them to move along a common scale. Low vocal and instrumental output must likewise be interpreted alongside freezing, rather than treated as evidence of safety or recovery.

Such dissociation is already important within the defensive repertoire. The Fear Threshold Model proposes that overt defensive reactions and putative nonverbal indicators of subjective fear need not change at the same level of threat. Developed from a systematic review and meta-analysis (11), the model received empirical support from differences between freezing and 22-kHz calling across shock intensity, cue timing, and anxiety-related variation (12). Calling could emerge later while freezing remained comparatively stable. This evidence motivates a differentiated account of fear-related behavior, but leaves a further question open: how do these defensive patterns relate to vocal engagement and instrumental action as the situation becomes less threatening?

Following the animal through extinction brings that question into focus. Extinction can involve new inhibitory learning and context-dependent retrieval rather than erasure of the original threat memory; defensive responding can subsequently return with time or a change of context (13–17). A reduction during extinction and its durability at a later test are thus separate questions. Neither, on its own, establishes whether the animal has resumed activities disrupted by threat. For preclinical models relevant to exposure-based treatment, the additional functional question is whether changes in defense are accompanied by renewed engagement and task performance, and whether those changes persist. Recovery in this functional sense need not recreate an earlier response pattern. Learning may establish a different organization of engagement and defense, whose appropriateness depends on the circumstances. This is a question about an individual’s history as well as its current response. The same low level of freezing can have different functional implications before aversive learning, during extinction, and after a relapse challenge.

Tri-reference-point (TRP) theory offers a framework for relating these possibilities to the demands of the situation. Originally formulated for decision making under risk, it distinguishes a minimum requirement, the status quo, and a goal, defining regions of failure, loss, gain, and success (18). Established accounts of threat imminence, risk assessment, and foraging under predation risk explain how danger changes the available behavioral priorities (19–22). We extend TRP’s reference-dependent logic to affective organization because avoiding unacceptable outcomes and pursuing attainable ones both require the prioritization of action. The extension relates protection from danger and engagement with opportunities to learned reference conditions (Figure 1A). A response combination acquires functional meaning from those conditions; low output is not itself a reference point, and pressing need not indicate that a goal has been attained. The Fear Threshold Model motivates differentiation on the danger-facing side, while the broader framework asks what replaces defense. This proposal organizes testable questions without requiring a literal psychological axis or assuming that rats implement TRP as a decision algorithm.

**Figure 1.**
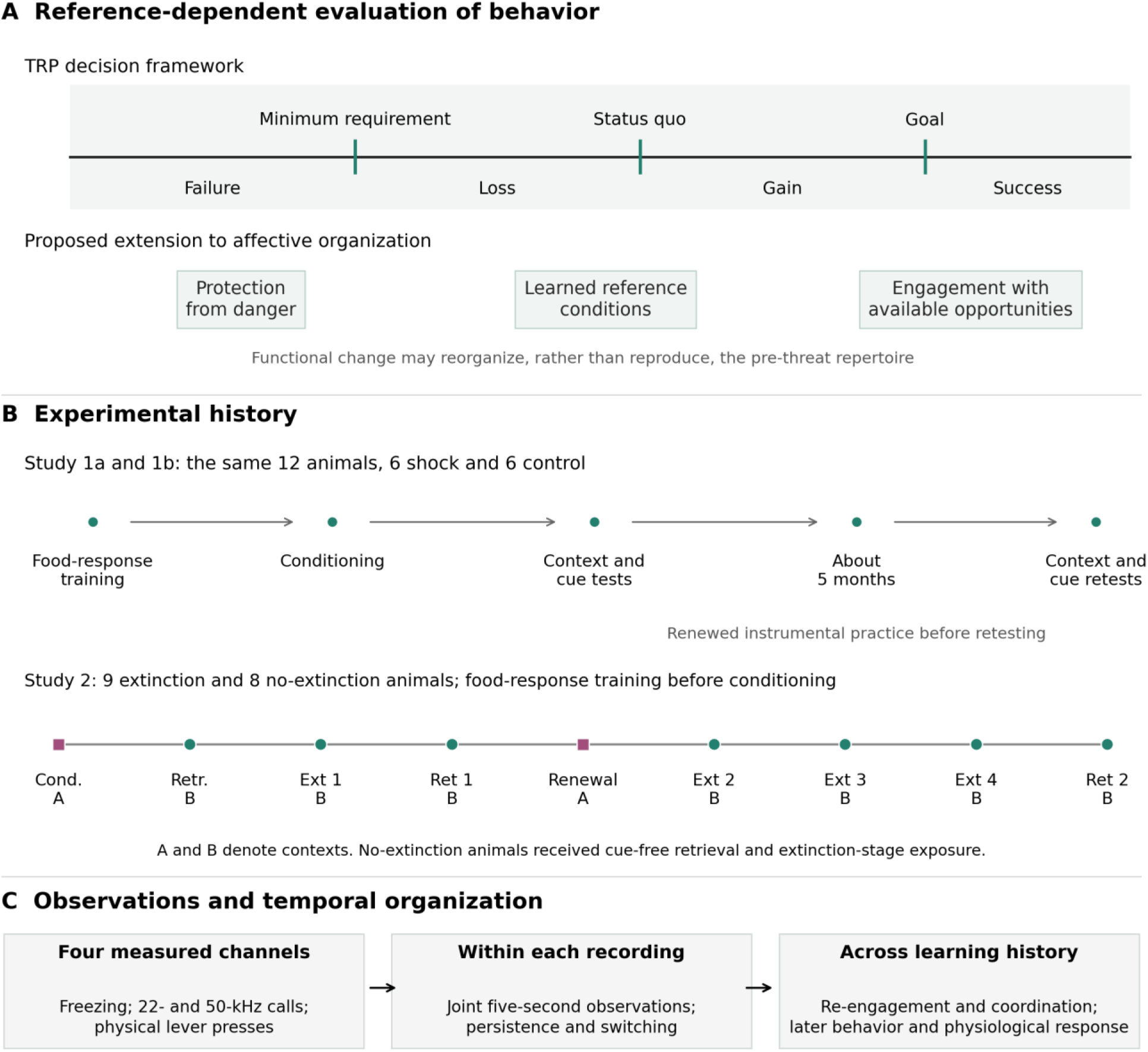
A functional question across threat learning and subsequent experience. (A) Tri-reference-point theory distinguishes a minimum requirement, the status quo and a goal in decision making. The proposed affective extension relates protection from danger and engagement with available opportunities to learned reference conditions. The diagram does not assign low output to a TRP region, estimate the reference points, or impose psychological identities on fitted components. Functional change may reorganize rather than reproduce the pre-threat repertoire. (B) Experimental histories. Studies 1a and 1b followed the same 12 animals, six shocked and six unshocked controls, through conditioning, cue and context tests, and retesting approximately five months later. Renewed instrumental practice preceded delayed testing. Study 2 followed nine extinction and eight no-extinction animals through conditioning, retrieval, extinction or matched context exposure, retention and renewal tests, further extinction or exposure, and final retention. A and B identify chamber contexts. Cond., conditioning; Retr., retrieval; Ext, extinction-stage session; Ret, retention test. Stage spacing is schematic, not elapsed time. Exact cohort-specific schedules are in Table S1. (C) Freezing, ultrasonic calls and physical lever presses provide complementary observations. Temporal models describe within-recording organization; experimental history defines comparisons between recordings. Fecal boli provide a separate acquisition-level physiological outcome. In Study 2, magenta squares mark context A and green circles context B.

To connect this framework to behavior, the analysis must retain both the different channels and their temporal organization. Computational neuroethology emphasizes structured behavioral sequences (23–25), and behavior-systems modeling in experimental psychology similarly uses hidden states to describe the organization of motivated action (26). A hidden Markov model (HMM) provides one way to examine whether recurring combinations of freezing, calls, and presses persist and give way to other combinations (27, 28). The observations remain the measured behaviors; the inferred states summarize patterns in their joint distribution, while transitions describe dependence between successive windows. Response intensity can change within a configuration as well as through switching between configurations. This distinction also separates two timescales. The HMM describes seconds-scale organization within a recording. Conditioning, extinction, elapsed time, and context define the learned histories across which those recordings are compared; they are not memory states estimated by the HMM.

The biological interpretation of an inferred configuration consequently depends on its measured signature and experimental context. Functional accounts of emotion use coordinated behavioral and physiological responses, including their persistence and generalization, to reason about internal states (29). Here, such reasoning does not turn a statistical state into a direct measure of subjective feeling. Freezing and vocalization are both objectively recorded, although their proposed inferential roles differ: freezing indexes a defensive reaction, whereas a 22-kHz call may also carry information relevant to subjective fear alongside its communicative and adaptive functions (11, 12). More broadly, the relationship between observed defenses and subjective experience remains debated (30–32). The distinction made here is between the evidence supplied by different responses and the experiences inferred from them, not between conscious and non-conscious behavior. It leaves open an experimentally tractable question about how these responses are organized when defense and reward pursuit compete.

We address that question in related rat studies spanning aversive conditioning, extinction, delayed retesting, and changes of context (Figure 1B,C; Table S1). Rats learned to press for food before threat was introduced, allowing later behavior to be interpreted against an established instrumental repertoire. Group-resolved comparisons establish how each measured channel changes across the experimental conditions. Temporal models then ask what their joint organization adds to those observations: whether recurring configurations are predictively useful, how their composition depends on modeling resolution and emission assumptions, and how their expression varies with learning history. The original four-region proposal motivates a coarse four-component comparison, while alternative resolutions test whether it adequately describes the observations. Associations with later behavior and physiological outcomes are evaluated separately from the prediction of behavioral sequences. The central aim is to characterize how established activity is suppressed and subsequently expressed in relation to defense, creating a basis for testing context-appropriate re-engagement rather than requiring return to one prescribed state.

## Results

### Defense and instrumental engagement across threat and time

We first asked whether changes in defensive immobility were accompanied by changes in a learned food-reinforced action. Study 1 compared six shock and six control animals through conditioning, cue generalization, and delayed retesting of the same animals (Figure 2; Table S2). At the first 9-kHz cue test, the shock group averaged 59.2% freezing, compared with 20.0% in controls. Physical lever-press rates were 1.13 and 3.47 per minute, respectively; one shock animal lacked a press record for this test. The separation in freezing was less pronounced at later cue tests, while instrumental responding remained variable across animals and conditions.

**Figure 2.**
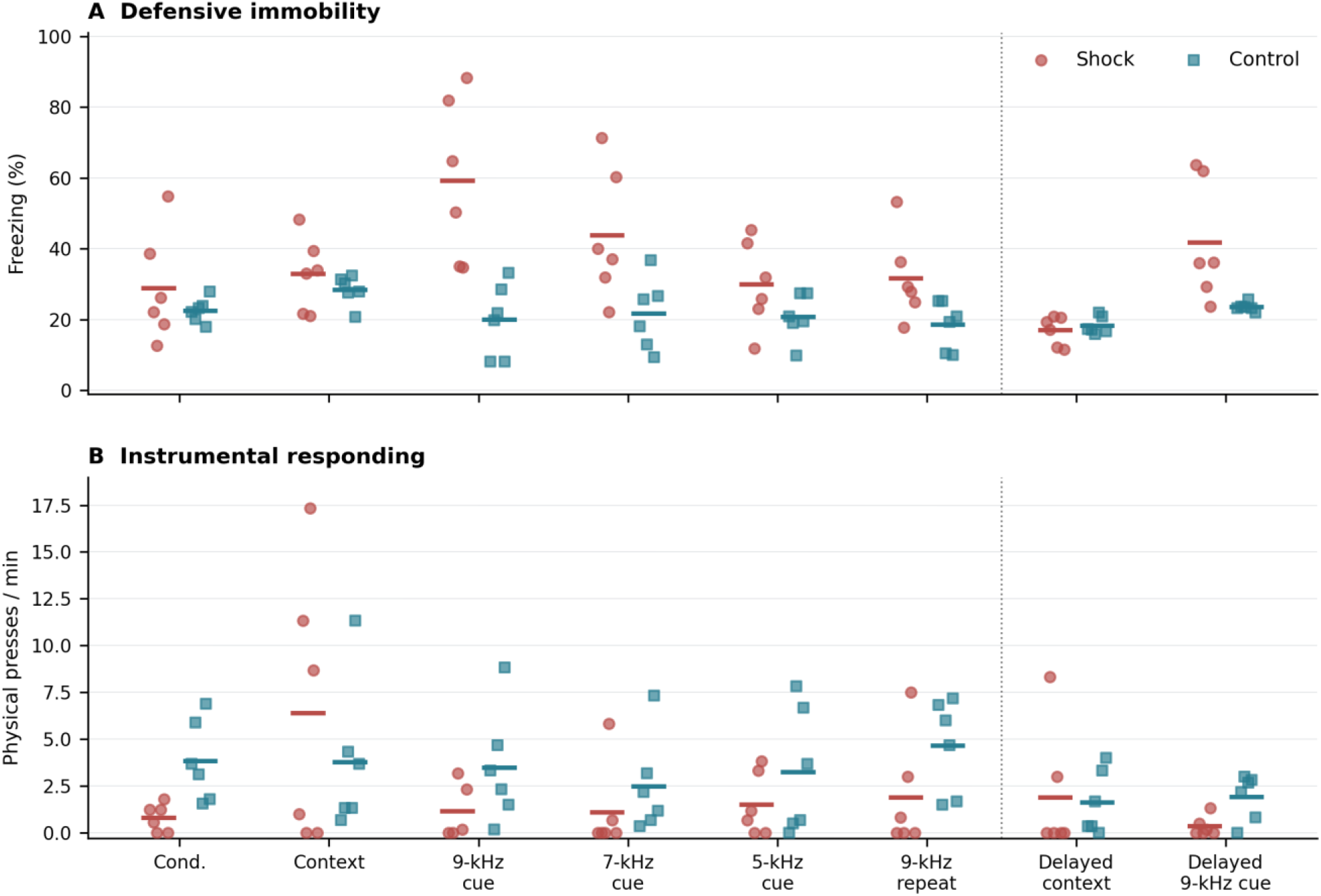
Defensive and instrumental behavior across initial learning and delayed tests. Freezing (A) and physical lever-press rate (B) in Study 1. Points are individual animals; short horizontal bars are group means. Each phase includes six shock and six control animals except first-cue pressing, for which one shock animal’s controller record was unavailable. The dotted divider separates initial testing from the approximately five-month retest of the same animals. Original acquisitions are shown; repeat acquisitions were not substituted. Conditioning uses the 600-1140-s task window, context tests use 0-180 s, and cue tests use 180-540 s, including the intervals between cues. Press rates use recorded physical inputs, the session-specific manual-end clock estimate and the actual covered duration. The 28 group comparisons included task-window levels and baseline-adjusted contrasts; none met the Holm-corrected threshold. The later observations combine elapsed time with intervening testing and renewed instrumental practice, rather than isolating a causal effect of time.

The no-shock controls also inform the interpretation of vocal activity during conditioning. In the investigator’s original 50-kHz summaries, the shock and control groups averaged 10.2 and 10.7 annotated events, respectively, during conditioning; their baseline summaries averaged 64.2 and 73.7 events. The conditioning-period group difference did not meet the false-discovery-rate threshold in the 49-comparison workbook family (q = 1.000; Table S2D). The summaries concern different windows and are not exposure-standardized rates. They do not resolve whether changes in calling reflect conditioning, habituation, or another feature of the experimental situation, and calling alone does not establish exploration.

The delayed tests showed why that variation matters for interpreting functional change. Approximately five months later, mean freezing in the shock group was lower during the context test than at the original test, decreasing from 32.9% to 17.0%, but mean pressing also decreased, from 6.39 to 1.89 per minute (Figure S1; Table S3). Control animals likewise showed lower means on both measures. Cue responses told a different story: delayed freezing in shock animals averaged 41.8%, below the first cue test’s 59.2% but above the repeated cue test’s 31.6%. Thus the impression of improvement depended on both the behavior measured and the earlier experience used as the comparison. These changes cannot be attributed to elapsed time alone, because the animals also differed in their intervening testing history and later experimental conditions.

The first-cue contrasts were sizeable on the observed scales: 39.2 percentage points more freezing and 2.34 fewer presses per minute in shock animals. The freezing comparison included six animals per group; the press comparison included five shock animals and six controls. The freezing contrast had an exact permutation p of 0.0022 and a Holm-adjusted p of 0.061 across the 28 group comparisons. None of the group or paired contrasts met its respective familywise threshold, but this does not turn the observed differences into evidence of absent learning or equivalent groups. Small groups, shared housing and repeated testing limit estimates of the magnitude and generality of these changes. The individual trajectories make the functional question concrete: lower freezing need not be accompanied by a higher press rate. Extinction provided a separate setting in which to follow these outcomes together.

### Behavior during extinction and later tests

We next followed defense and instrumental engagement through extinction and later tests. Freezing and physical lever presses were available for all 17 Study 2 animals across nine experimental stages, so these group comparisons did not depend on the availability of the other channels (Figure 3; Table S4).

**Figure 3.**
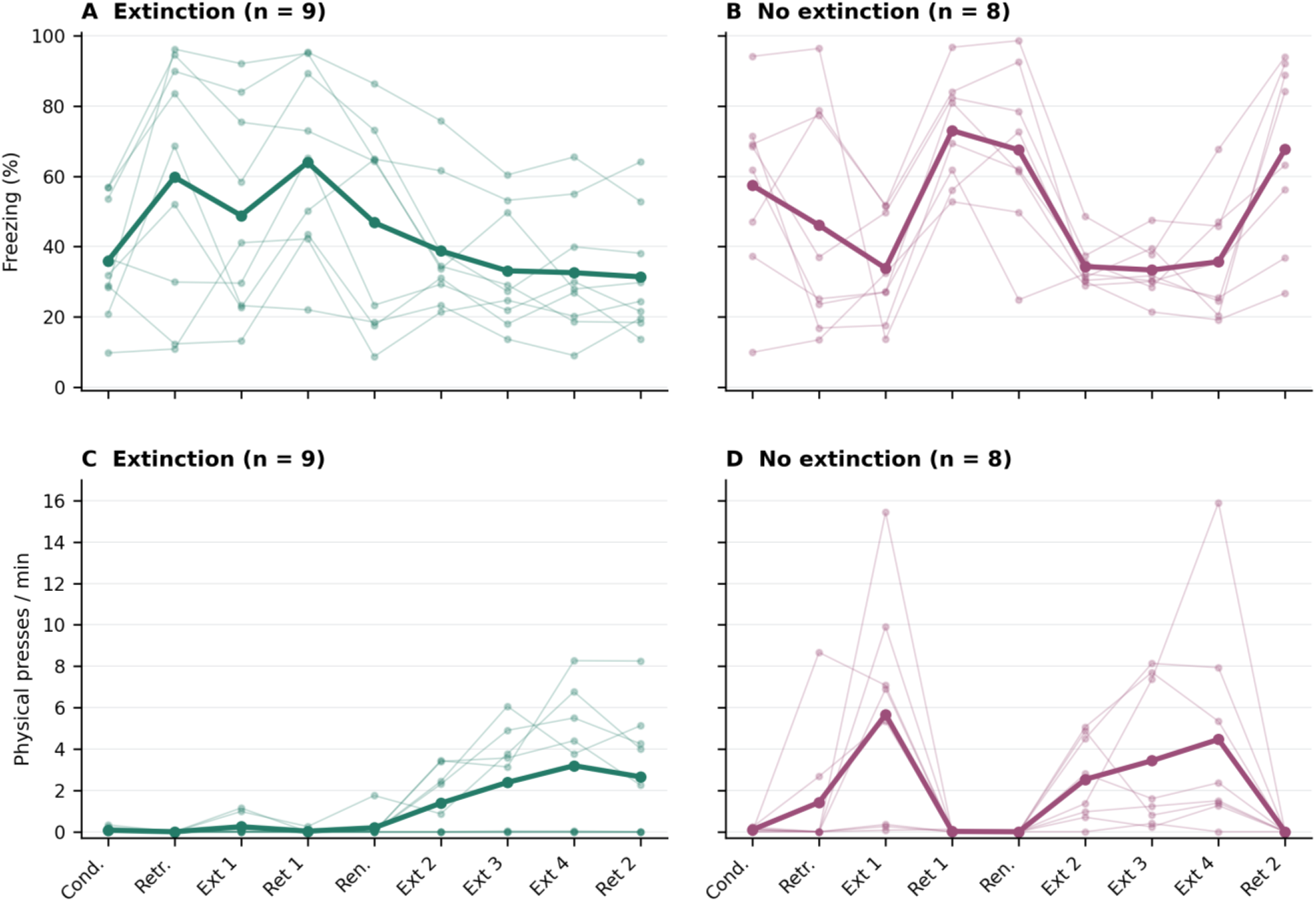
Observed defense and instrumental engagement during extinction and later tests. Task-window freezing (A, B) and physical lever-press rates (C, D) for all nine extinction and eight no-extinction animals in Study 2. Thin lines connect observations from the same animal; thick lines connect group means. All 17 animals contribute to every phase in these channel-specific analyses, independently of vocal availability or joint-model coverage. Task windows follow the executed acquisition schedules in Table S1. Rates use physical input counts divided by recorded task-window duration. Cond., conditioning; Retr., retrieval; Ext 1-4, extinction sessions in the extinction group and matched cue-free context exposure in the no-extinction group; Ret 1/2, retention tests; Ren., renewal. Phase spacing is categorical, not elapsed time. The descriptive trajectories do not establish functional recovery: none of the 36 planned between-group comparisons met the Holm-corrected threshold.

Before extinction began, conditioning-period freezing already differed between the future extinction and no-extinction groups (35.9% and 57.4%, respectively). The later contrasts must therefore be interpreted alongside these earlier differences and the cohort-specific schedules. In the nine extinction animals, mean task-window freezing was 48.8% in the first extinction session and 32.5% in the fourth, while pressing increased from 0.24 to 3.19 presses per minute. These descriptive changes capture two aspects of the animal’s response to its situation: the expression of defense and engagement in an already learned instrumental activity.

The course within a session was less uniform. During the first extinction session, the same animals averaged 44.0% freezing in the first 180 seconds of the task window and 45.2% in the last 180 seconds; pressing averaged 0.41 and 0 presses per minute, respectively (Figure S2; Table S5). The end of this session therefore did not show either uniformly low freezing or renewed pressing. Changes varied across animals and sessions, and none of the paired early-to-late comparisons met the familywise threshold. At the second retention test, the extinction group averaged 31.4% freezing and 2.65 presses per minute, compared with 67.8% freezing and no presses in the no-extinction group. The estimated extinction-minus-no-extinction differences were-36.4 percentage points and 2.65 presses per minute. These magnitudes are compatible with less defensive immobility and greater instrumental engagement in the extinction group, but the small samples and earlier group differences prevent attributing the entire contrast to extinction. The 36-comparison family gave Holm-adjusted p = 0.221 and 0.909, respectively.

Vocal activity added information beyond freezing and pressing. Investigator-scored task summaries were available for all 17 Study 2 animals across the nine phases, including recordings without sufficient joint coverage for the temporal model (Figure S3; Table S6). At the fourth extinction session, the summaries recorded no 22-kHz output in either group, although mean freezing in the extinction group remained 32.5%. At the second retention test, mean 22-kHz duration was 4.45 seconds in the extinction group and 126.37 seconds in the no-extinction group; mean annotated 50-kHz event counts were 30.6 and 41.2, respectively. These correspond to extinction-minus-no-extinction differences of-121.9 seconds and-10.7 annotated events, rather than a common shift on one affective scale. None of the 18 vocal group comparisons met the false-discovery-rate threshold; the post hoc within-cohort permutation sensitivity (minimum adjusted q = 0.066) likewise left their generality uncertain rather than showing no vocal response. These duration and count summaries retain their distinct units and task windows. Together, they distinguish the course of calling from that of defensive immobility.

### Temporal organization of the joint repertoire

Observing these channels separately leaves a further question: does their joint expression carry information that is lost when successive observations are treated independently? We examined 45,300 five-second windows from 29 animals (Table S1C,D) using a hidden Markov model and an independent mixture with the same four-component emission specification. Both vocal-source policies were examined on the same windows. All recordings from an animal remained together in five-fold cross-validation. The temporal model scored better for every animal, with a mean paired advantage of 0.468 nats per window under investigator-summary priority and 0.401 under retained-annotation priority. Within-recording temporal dependence therefore contributed predictive information under both observation policies (Figure 4A).

**Figure 4.**
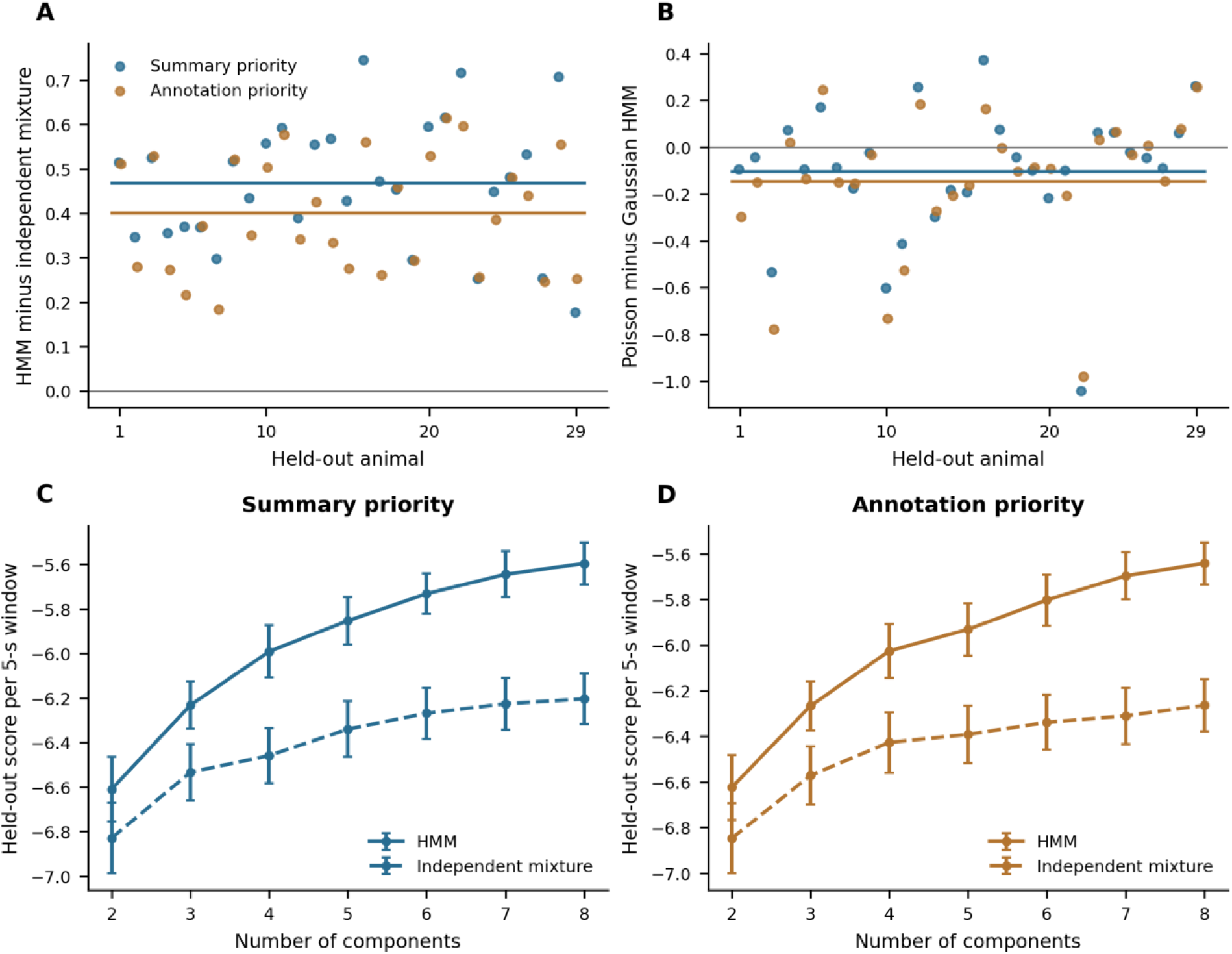
Temporal dependence and resolution shape predictive performance. (A) Paired held-out score differences between four-component Gaussian HMMs and matched independent mixtures at variance floor 0.1. Each point is one of 29 animals; horizontal lines are animal means. (B) Poisson-count minus Gaussian count-mass HMM scores on those same observations and folds, with the freezing floor matched at 0.1. (C, D) Animal-averaged held-out scores for two through eight components. Error bars are descriptive standard errors across animals, not independent validation-set confidence intervals. Higher scores are better within a source policy. Integer count channels contribute probability masses and freezing contributes a density in percentage units. Summary and annotation priority refer to alternative policies for discordant vocal sources, not independent datasets. Animal IDs are ordered consistently within the paired comparisons. All observations from an animal remained in one fold, including both Study 1 series. Component-count comparison is not nested selection or evidence for a unique natural number of states.

That advantage did not imply that four components exhaust the repertoire. Among models with two to eight components at the reference variance floor of 0.1, eight gave the highest animal-averaged held-out score under both source policies. Its paired gain over four components was 0.395 nats per window under summary priority and 0.384 under annotation priority, with a positive gain for every animal. Seven and eight fell within one descriptive standard error of the best HMM score (Figure 4C,D). We therefore examined four components as the coarse scale motivated by the proposed functional regions, alongside the finer eight-component description (Figure S4). The search ended at eight, and model resolution was evaluated using the same cross-validation scores. It identifies useful resolutions within the tested range rather than a unique natural state count or an independently validated optimum.

### Low output includes distinct freezing configurations

At the four-component scale, two configurations contained very little vocal or instrumental output but differed markedly in freezing: 15.8% in one and 74.0% in the other (Figure 5A). A vocal-active configuration combined 22-kHz and 50-kHz calls with substantial freezing (52.9%) and almost no pressing. A press-active configuration averaged 1.88 registered inputs per five seconds with relatively little freezing (14.8%). It contained 99.4% of all registered inputs; the corresponding annotation-priority configuration averaged 1.97 inputs and 14.9% freezing and also contained 99.4% of inputs. Lever inputs contributed to model fitting; this percentage describes their concentration in the fitted partition, not accuracy against independently known behavioral states. Physical pressing therefore differentiated instrumental engagement sharply in both selected descriptions. Defensive immobility could occur with or without frequent calls, while instrumental activity distinguished another part of the repertoire. The pooled occupancies of the lower-freezing low-output, higher-freezing low-output, vocal-active and press-active configurations were 45.8%, 27.1%, 15.1%, 11.9%, respectively. These proportions reflect the phases and recording durations contributed by the design, not the proportions expected in a healthy or recovered animal.

**Figure 5.**
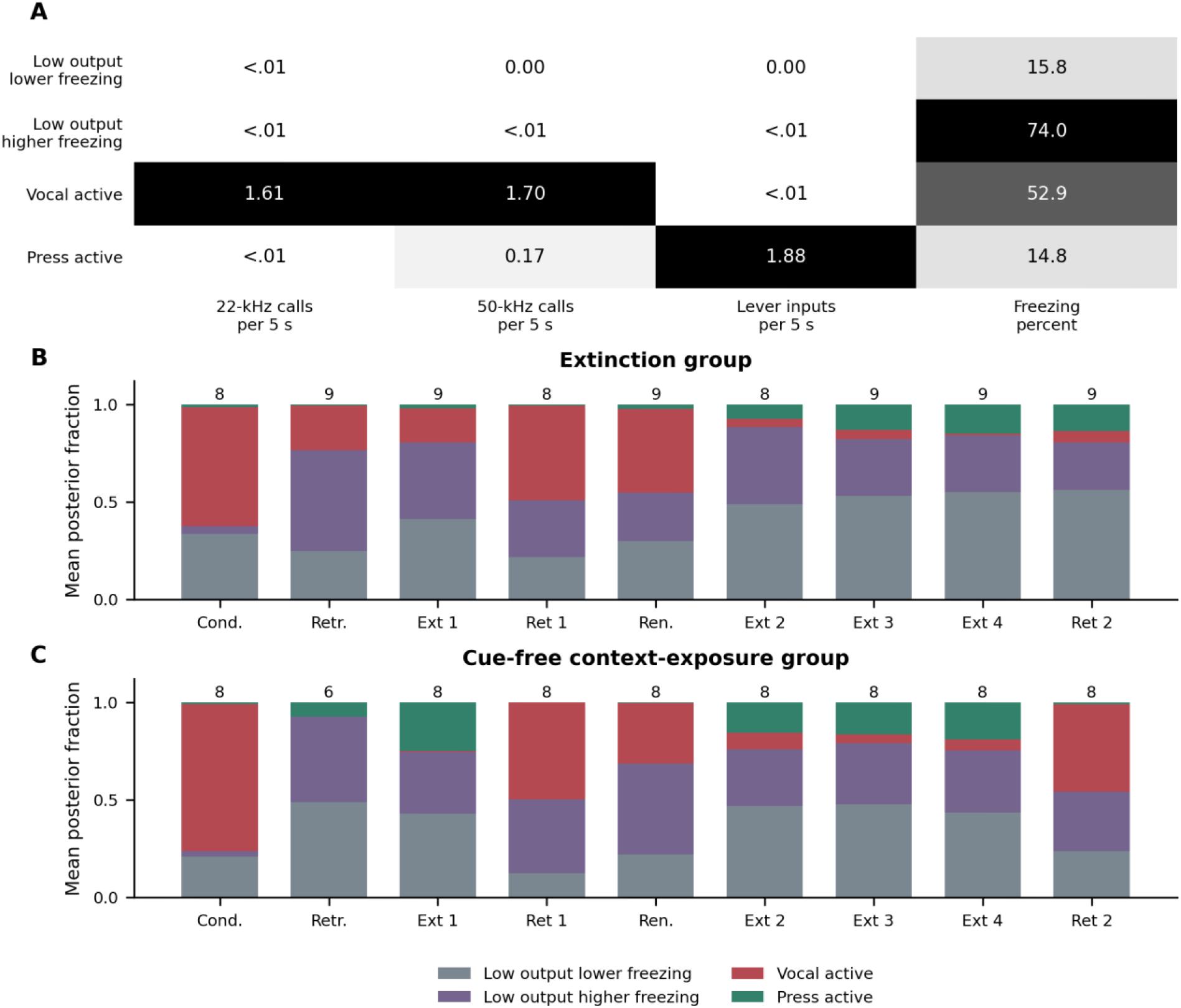
Low vocal and instrumental output accompanies distinct freezing configurations. (A) Empirical channel means among mass-decoded windows of the selected K=4 Gaussian HMM, variance floor 0.1, investigator-summary priority: 45,300 windows from 29 animals and 208 acquisitions. Cell shading is scaled within channel; cell values retain their own units. Values between zero and 0.01 event per window are shown as <.01. (B, C) Task-window posterior fractions averaged equally across included animals. At least 95% joint task coverage is required; numbers above bars are animals, not independent windows. Cue-free context exposures are included under the declared approximate timing policy. Missing coverage is not zero occupancy. Cond., conditioning; Retr., retrieval; Ext 1-4, extinction or matched context exposure; Ret 1/2, retention; Ren., renewal. Colors describe measured configurations rather than established affective states. Changes in sample composition and the learned histories contributed by the design prevent pooled fractions from being read as healthy or recovered targets.

The contrast between low output and low freezing is central to interpreting these patterns. A quiet interval may contain little or substantial freezing; the absence of calls and presses does not distinguish those conditions. The two selected source-priority fits gave 98.8% agreement after label alignment, but agreement across all valid starts was less consistent: median pairwise normalized mutual information (1 denotes identical partitions) was 0.613 under summary priority and 0.914 under annotation priority. The selected configurations are therefore useful descriptions, not a uniquely recovered four-part architecture. At eight components, low-output configurations in the summary-priority fit averaged 10.2%, 48.2% and 94.6% freezing. Other components separated 22-kHz-dominant, 50-kHz-dominant, mixed-vocal and instrumental patterns (Figure S4). Finer resolution adds distinctions without establishing a common sequence of affective stages or a simple refinement of every four-component state.

We then asked whether response combinations persisted when calls and presses were modeled directly as counts, while retaining temporal dependence. At four components and the same 0.1 freezing-variance floor, Poisson-count HMMs scored lower on average than their Gaussian count-mass comparators: by 0.103 nats per window under summary priority and 0.145 under annotation priority (Figure 4B). The count models nevertheless gave different partitions, with aligned agreement of 68.0% and 52.1%, respectively, with the Gaussian fits. They retained an instrumental-rich, lower-freezing configuration, but grouped pressing with more quiet intervals while dividing vocal activity differently (Figure S5). Instrumental engagement remained recognizable even though its configuration boundaries changed. The Gaussian specification performed better in this matched predictive comparison.

These configurations differed not only in composition but also in how they persisted and gave way to one another. In the selected summary-priority K=4 model, fitted probabilities of remaining in the same configuration at the next five-second step were 0.766 for lower-freezing low output, 0.810 for higher-freezing low output, 0.824 for vocal activity, and 0.497 for press activity (Table S7). The press-active pattern was therefore represented as more transient than the other three at this resolution. The complete matrices retain fitted probabilities and observed adjacent counts for persistence and every off-diagonal transition, pairing only contiguous observations within an acquisition. Within these selected representations, instrumental activity formed shorter bouts among more persistent vocal and low-output configurations.

### Expression across learning and later tests

The expression of these configurations also differed across the original conditioning groups. In the summary-priority four-component model, mean posterior occupancy of the vocal-active configuration increased from 18.1% during baseline to 53.0% during conditioning in shocked animals, but decreased from 22.1% to 4.0% in controls (six animals per group; Table S2E). Press-active occupancy changed from 29.9% to 5.3% in shocked animals and from 24.7% to 22.4% in controls. The joint description captures both suppressed instrumental expression and increased vocal activity in the shocked animals.

The Study 2 recordings extended this account across different histories. Vocal-active behavior occupied much of conditioning in both groups, whereas retrieval in extinction animals included more high-freezing, low-output behavior. At the first retention test, the vocal-active and high-freezing low-output configurations together occupied most of the task window in both groups (Figure 5B,C). All nine extinction animals contributed to the first-to-fourth-extinction descriptions. Their mean press-active posterior fraction increased from 2.0% to 15.1%, while the vocal-active fraction decreased from 17.5% to 0.8%. Lower-freezing low output increased from 41.3% to 55.1%, and higher-freezing low output decreased from 39.2% to 29.0%. These descriptive changes show renewed instrumental expression alongside continued defensive immobility, rather than a uniform passage into one recovered configuration.

These trajectories also depend on which animals and intervals can be jointly observed. At least 95% task coverage was available in 148 of the 153 Study 2 acquisitions, including every animal at the final retention test and all 32 cue-free context-exposure acquisitions. Cue-free recordings remained represented under an approximate audio-clock policy. Feeder-linked relative clocks gave a similar selected four-component partition on common windows, while audio-to-video timing remained uncertain. In Study 1, alternative lever origins changed about 20% of four-component assignments, whereas the notebook-based audio alternative changed less than 1% (Figure S6). The measured group trajectories and the timing-dependent joint configurations therefore provide complementary descriptions.

### Post-extinction behavior and later responding

Whether a behavior returns at the group level is distinct from whether an individual’s behavior anticipates its later response. In the nine extinction animals, late pressing in the fourth extinction session was positively related to pressing at the second retention test (Spearman rho = 0.877; exact cohort-restricted permutation p = 0.0083). The association did not meet the familywise threshold across the 24 specified follow-up comparisons (Holm-adjusted p = 0.200; Table S8), and none of the other associations did so either. Four comparisons were not estimable because a measure was constant; sparse pressing and tied values limited several others. The observed relationship motivates testing continuity of instrumental engagement in a larger longitudinal sample, but does not yet support individual prognosis or a claim that post-extinction behavior predicts relapse.

Relating inferred configurations to later behavior requires joint observations during extinction, not merely complete records of the later response. All nine extinction animals contributed final-180-second configuration fractions for the three chronological pairs (Table S9). Press-active occupancy in the fourth extinction session was positively related to second-retention pressing under both source policies (rho = 0.923; exact cohort-restricted permutation p = 0.0028; Holm-adjusted p = 0.0667). The two policies describe the same animals, not independent replications. No configuration association met its 24-comparison familywise threshold. This also held for whole-task and hard fractions, eight-component descriptions and the alternative clocks. The press-active correspondence retains the continuity visible in raw pressing, but pressing also contributes to the configuration and the representation uses the full behavioral dataset. Its additional predictive value therefore requires the animal-excluded comparisons below.

### Behavioral organization and physiological response

The final question was whether the inferred configurations carried information beyond the behavioral channels used to construct them. Fecal boli, counted after each acquisition, supplied an external physiological outcome. Across all 153 Study 2 acquisitions from 17 animals, mean freezing was positively associated with boli count after adjustment for phase, group and learning cohort (negative-binomial GEE, coefficient per 10 percentage points = 0.115, 95% CI 0.005 to 0.224). With recording duration included as an exposure offset, the estimate remained positive but its interval included zero (coefficient = 0.100, 95% CI-0.006 to 0.206; Figure 6; Table S10). This is limited evidence of a whole-session association, not of a physiological change accompanying each inferred transition.

**Figure 6.**
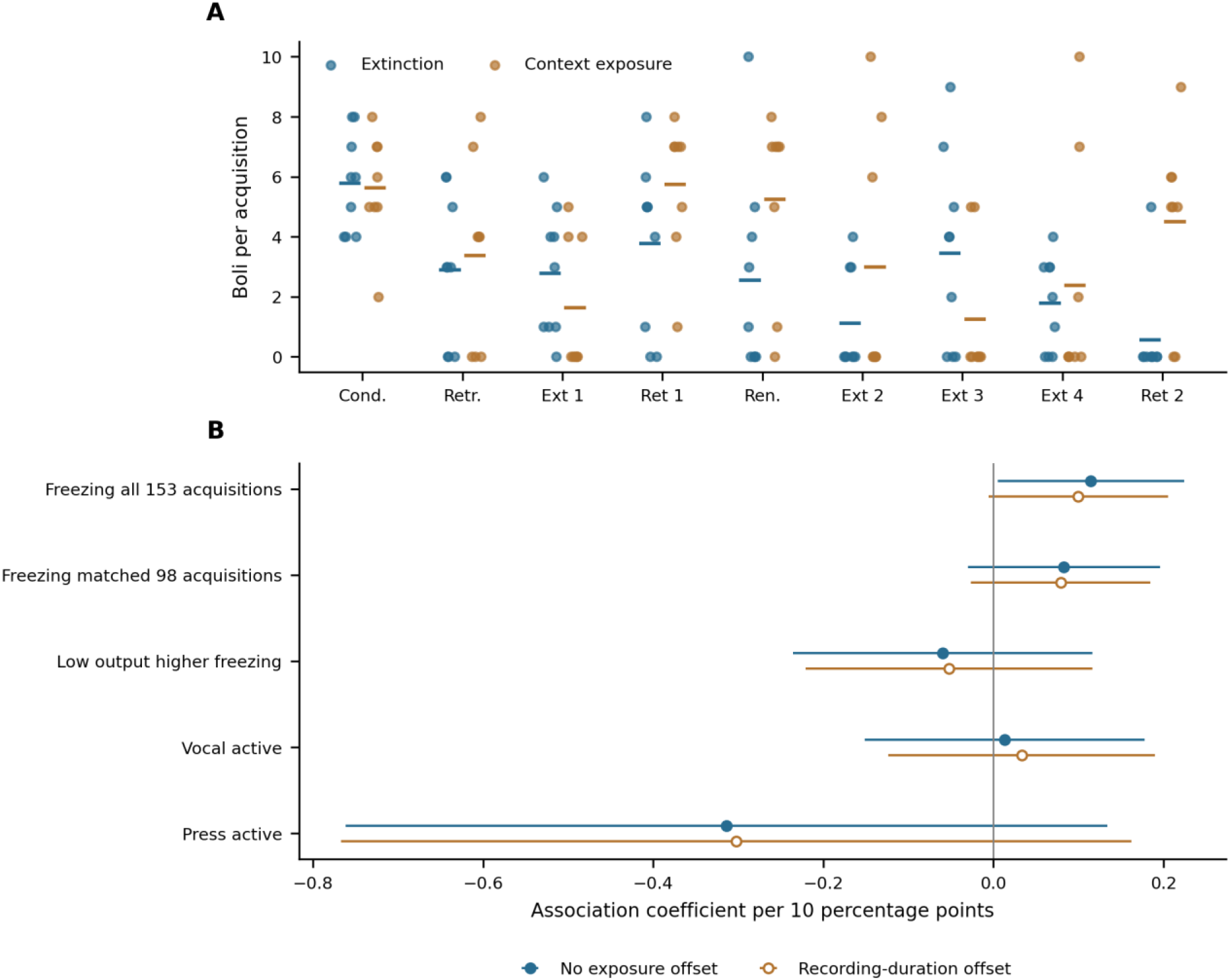
Acquisition-level physiological associations. (A) Fecal-boli counts in all 153 Study 2 acquisitions from 17 animals. Points are individual observations; horizontal bars show group means and jitter separates overlapping points. (B) Negative-binomial GEE coefficients and unadjusted, bias-reduced 95% confidence intervals with animal clusters and adjustment for phase, group and learning cohort. Filled symbols omit an exposure offset; open symbols add log recorded duration in minutes. The first freezing estimate uses all 153 acquisitions, while the matched freezing and displayed configuration estimates use 98 acquisitions with 95% whole-recording joint coverage, still representing 17 animals. Configuration contrasts represent a ten-percentage-point redistribution from the lower-freezing low-output reference while holding other proportions constant. None of the 48 configuration coefficients across source, clock, window and fraction definitions met the Holm threshold, with or without the duration sensitivity. The task-window models relate 148 task observations to whole-acquisition boli totals. Different significance outcomes between models or observation scopes do not establish superiority.

Whole-acquisition configuration analysis included 98 acquisitions from the same 17 animals; task-window analyses included 148 acquisitions. No occupancy coefficient met the Holm threshold across the 48 source-policy, clock, window and fraction-definition comparisons, with or without the recording-duration sensitivity. The matched-subset freezing estimates were also imprecise. These comparisons do not establish that configurations add physiological information beyond freezing, nor does a significant coefficient in a larger sample establish superiority over a nonsignificant coefficient in a smaller one.

These association models identify neither the time of defecation nor added prediction in animals excluded from model fitting. The latter requires a matched held-out comparison against the underlying behavioral measures.

### Added prediction of later and physiological outcomes

To test added prediction directly, each target animal was excluded from both its HMM representation and the outcome model. In the nine extinction animals, configuration fractions from the final 180 seconds were added to raw antecedent behavior to forecast later freezing or pressing. Only three of 48 primary comparisons reduced mean squared error, all for fourth extinction to the second retention test. Their augmented freezing RMSEs were 19.17, 14.66 and 17.91 percentage points, each above the 12.38-point error of simply carrying forward late-extinction freezing. Their average advantage over their fitted raw baseline also changed sign when the largest benefiting animal’s loss was omitted. The complete 144-comparison sensitivity retained both sources, K=4 and K=8, two raw baselines and three fixed ridge penalties (Table S11). This is exploratory internal validation, not evidence of independent individual prognosis.

For fecal boli, the primary held-out comparisons used the same 98 whole acquisitions from all 17 animals, with phase, group, cohort and recording-duration adjustment. Adding configuration fractions to freezing reduced average held-out log score by 0.034 and 0.033 per acquisition at K=4, and by 0.091 and 0.104 at K=8, under summary and annotation priority, respectively. Scores were averaged within animal before averaging across animals. None of the 32 source, resolution, raw-baseline, window and offset comparisons improved average log score (Table S12). These models therefore supplied no consistent added outcome-prediction benefit under the tested specifications. That finding concerns the available acquisition-level physiological outcome, not the validity of the behavioral sequences or every possible physiological correlate.

## Discussion

Following recovery from threat requires knowing what replaces defensive behavior. In these experiments, lower freezing was not consistently accompanied by greater instrumental engagement, and low vocal and instrumental output occurred at both low and high levels of freezing. Physical inputs also identified a clearly differentiated instrumental-active configuration. The temporal models added a complementary result: recent behavioral sequence improved prediction of subsequent observations beyond a matched independent mixture. Together, these findings make recovery a question about the composition and temporal organization of engagement, not simply the disappearance of defense. Familiar responses may be coordinated differently after aversive experience. The present evidence characterizes that reorganization without establishing four distinct affective modes or demonstrating that a new adaptive state was learned.

The experimental sequence makes that question tangible. Rats had learned an effective food-reinforced action before aversive experience changed the circumstances in which it was performed. At the delayed context test, lower group-mean freezing accompanied lower, not higher, pressing. Across extinction sessions in a separate cohort, the descriptive pattern was different: freezing decreased while pressing increased. Some observed contrasts were sizeable, but they do not isolate contrasting recovery mechanisms. Small groups, pre-extinction differences and cohort-specific procedures limit their precision and generality; an adjusted threshold does not decide their biological importance or establish that a manipulation was ineffective. They show why the two outcomes must nevertheless be measured separately. A later test can appear improved with respect to defense while showing no return of instrumental engagement; conversely, resumption of that engagement need not wait for defense to disappear. Earlier behavior provides a reference for this change, not a requirement to reproduce the same response frequencies or sequence. Functional evaluation must also ask whether the resulting engagement fits the available opportunities and remaining danger. A pooled proportion of time assigned to a component cannot answer that question.

The multimodal configurations extend this distinction within the defensive repertoire itself. High freezing appeared both with little calling and with substantial vocal output. At finer resolution, both call bands contributed configurations with different freezing levels, while instrumental responding occupied another part of the repertoire. A scalar account that orders all responses from less to more fear would lose these differences. This result is compatible with the Fear Threshold Model’s proposal that defensive reactions and vocal indicators can be recruited differently (11, 12), but it is not a new measurement of their thresholds. Establishing recruitment order would require graded, experimentally controlled threat and sufficiently resolved observations of each response. Subdividing a continuous freezing measure into several HMM components cannot provide that evidence by itself. Nor do the present vocal configurations decide whether 50-kHz calls reflect exploration, anticipation, social communication, or another context-dependent process (7–10).

These findings clarify what TRP contributes to the analysis. Its original reference points concern decision making under risk (18); their application here organizes questions about protection from danger, the learned current condition and engagement with available opportunities. The experimental sequence and multimodal measurements were motivated by this functional extension, but empirical configurations need not correspond one-to-one to its regions. Low output cannot identify a status quo or a safe condition, and instrumental and vocal engagement need not occupy separate states. The proposed extension is therefore constrained by the measurements rather than confirmed by choosing four components. Changing danger while holding reward opportunity constant, or changing reward value while holding threat history constant, would test how behavior reorganizes relative to independently specified references. The present design does not isolate all three reference points or test their distinctive priority ordering. It does show why a functional account must follow the behavior accompanying defensive change.

Temporal modeling is useful for that program because behavior has an order as well as a composition. A freezing episode, a period of calling, and a sequence of presses are not interchangeable with the same observations randomly distributed through a recording. The predictive benefit of the HMM over its matched independent mixture supports this temporal dependence, in line with ethological and behavior-systems approaches to motivated action (23–26). Within the selected four-component Gaussian descriptions, press-active configurations were more transient than vocal-active or low-output configurations. Brief pauses need not indicate loss of task engagement: they can also reflect the programmed delay between a response and feeder activation. The predictive advantage does not show that a discrete model is superior to every continuous dynamical account: persistence, autocorrelation, and changing experimental conditions can all contribute to the advantage. The two relevant timescales must also remain separate. The HMM represents dependence between successive five-second windows; conditioning, extinction, delay, and context supply the histories across which those within-recording patterns are compared. The model does not infer a memory state that develops across sessions.

The number and identity of configurations are correspondingly empirical choices rather than theoretical givens. Finer Gaussian models predicted better than the four-component description within the tested range, and multiple initialization basins produced different partitions. Count-based HMMs also reorganized the response combinations, despite retaining temporal dependence. Their lower mean predictive scores in this comparison do not make the Gaussian labels a natural taxonomy. Four components retain value as a coarse comparison scale, but the meaningful biological question is which response relations and changes remain recognizable across defensible representations. Low output accompanied by different levels of freezing is one such distinction in the Gaussian descriptions. A functional change can also reorganize familiar responses without creating an additional statistical component. Because the present models share configurations across recordings, they do not estimate the learning process by which a new adaptive organization might emerge. Independent experiments must establish which distinctions improve intervention evaluation.

This behavioral level can inform neural experiments without substituting for them. Freezing itself can involve coordinated autonomic and central processes rather than a simple absence of action (33). Circuit studies describe how defensive responses are organized (34), including competitive interactions between defined central-amygdala cell populations that regulate conditioned flight and freezing (35). Accounts of threat-related decisions further emphasize how the available response repertoire changes with the spatial and temporal proximity of danger (36). Such work makes persistence and changes in behavioral configuration plausible targets for neural investigation. It does not identify the present HMM components with those circuits. A mechanistic test would ask whether a targeted manipulation changes particular response combinations or their persistence, and whether the effect survives comparison with the underlying movements, calls, and presses. No neural mechanism or dedicated transition gate is inferred from these recordings.

Physiological and later behavioral outcomes impose a separate test of functional relevance. Here the whole-session freezing association with fecal boli was sensitive to adjustment for recording duration, and occupancy associations were imprecise. The direct held-out analyses then found no consistent added outcome-prediction benefit from configuration summaries. Higher predictive resolution for behavioral sequences did not translate automatically into better prediction of later responding or boli counts. These exploratory comparisons used only 17 animals for physiological outcomes and nine for later behavior; adding configuration features also increases model complexity. Their results delimit the utility demonstrated by the specified pipelines, rather than establishing that all temporal summaries lack functional information. Larger independent longitudinal samples are needed to test whether an outcome benefit generalizes. Reproducibility and interpretable behavioral signatures cannot substitute for that empirical evaluation.

The translational implication is a design principle for preclinical evaluation, not a demonstrated recovery assay. Extinction research has established why reduced defensive expression during treatment and its persistence across time and context are different problems (13–17). Functional assessment adds another: whether the individual can engage in relevant activities under those conditions (32). Multimodal measurement supplies the observations, the TRP extension identifies the functional questions, and temporal modeling describes their organization. None alone measures wellbeing. An individual’s pre-threat repertoire is an informative reference, but adaptive change may require a different balance of engagement and defense. Returning indiscriminately to an old action or location is not necessarily beneficial. The relevant criterion is context-appropriate function that can be sustained or adjusted as circumstances change.

This perspective also gives comparative and experimental psychology a concrete role in the next tests. Food pressing is an observable learned action here, not proof of goal-directed control in the technical sense, which would require tests such as outcome devaluation or contingency degradation (37). Exploration should likewise be measured directly rather than inferred solely from pooled calls. Across species, the transferable question is how protective responses and engagement with opportunities are organized, not whether the same vocalizations, motor acts, or number of states recur. Species-appropriate measures of exploration, foraging, social engagement, and avoidance could test corresponding functional relations while allowing the statistical configurations to differ. That comparison would separate a general account of behavioral priorities from features specific to this task and its measurements.

The present scope is limited by small, male-only cohorts from one laboratory, shared housing, and the reuse of the first cohort for delayed testing. Later differences combine prior experience, elapsed time, and testing conditions, including renewed instrumental practice before delayed retesting, so they cannot isolate a causal effect of time. Study 2 animals also had prior experimental use that could not be fully characterized for adjustment. Joint recording coverage is narrower than the available single-channel evidence and varies across phases; timing alternatives alter some short-window assignments. The source and alignment comparisons therefore delimit the configurations without discarding otherwise observed freezing, pressing, or investigator-scored vocal activity. Model constraints, five-second aggregation, and the lack of retained call-subtype classifications further limit biological specificity. These limitations preclude a claim of validated individual prognosis, but do not remove the need to distinguish defense from the functions an animal may or may not resume.

A prospective test should follow that distinction from the outset: measure each animal’s repertoire before threat, track its reorganization during learning and intervention, and evaluate later behavior in both familiar and changed circumstances. The timing of sustained re-engagement, its intensity and persistence, and the responses accompanying it provide separate candidate endpoints. Models should be fixed before evaluation in independent animals and compared with simpler measures on the same observations. The central endpoint is whether the individual engages appropriately and can sustain or adapt that engagement, rather than whether it reproduces a prescribed state occupancy. This approach makes functional recovery experimentally testable as a possible reorganization of behavior, beyond defensive decline alone.

## Materials and methods

### Study design and animals

The study combined aversive learning with an established food-reinforced response to examine defense and instrumental engagement across changes in experience. Study 1a compared 12 male Sprague-Dawley rats allocated to shock and unshocked control groups, six per group. Study 1b retested the same animals approximately five months later. Study 2 began with 18 males assigned to extinction and no-extinction groups by alternating allocation after ranking open-field center occupancy. One no-extinction animal was excluded before conditioning after food refusal and abnormal behavior and was housed separately, leaving nine extinction and eight no-extinction animals. The animal, not a five-second observation or a repeated acquisition, was the experimental unit.

Study 1a animals weighed 300-350 g and were reported as approximately 6-8 months old; Study 2 animals weighed 350-400 g and were reported as approximately 12 months old. Rats were obtained from Beijing HFK Bioscience, China (SCXK (Beijing) 2019-0008; Sprague-Dawley, RRID:RGD_70508). They were housed three per cage with a reversed light cycle, lights on from 20:00 to 08:00, daily rationed food, and freely available water. Animals were handled for five minutes daily during the initial acclimation week, and body weight was monitored daily from the week before testing through the experimental period. The Study 2 animals had been used by an earlier experimenter; their complete preceding exposure histories were not available for adjustment, and they are not assumed to have been experimentally naive. Only males were studied.

The Fujian Normal University animal ethics committee approved Study 1a under IACUC-20230054, Study 1b under IACUC-20240014, and Study 2 under IACUC-20240070. The original Study 1 sample was chosen for exploratory within-animal comparisons under an animal-reduction rationale. No prospective registration or power analysis was performed. Study 1a allocation was described as random in the experimental protocol, but the allocation sequence was not retained. Study 2 allocation was balanced rather than a documented randomization procedure. The vocal-curation protocol specified assessors unaware of the study aims; masking to experimental group during collection, scoring, and computational analysis was not documented. Reporting uses ARRIVE 2.0 and STRANGE to identify the design and sampling information available for these experiments (38, 39).

### Apparatus and instrumental training

Conditioning and testing took place in two Coulbourn chambers measuring 31 x 26 x 33 cm, each fitted with a stainless-steel grid floor, speaker, chamber light, food dispenser, and retractable lever. The chambers were placed in sound-attenuating compartments, vertically separated with foam between them. Context A retained the grid floor and standard chamber walls. Context B used a raised acrylic floor wrapped in black tape, blue nonwoven wall coverings, and 5% acetic-acid odor. A collection tray beneath the floor allowed fecal boli to be counted after each acquisition. Chambers were cleaned between animals. These contextual manipulations changed several sensory features together rather than isolating one contextual cue.

Instrumental training preceded conditioning. On the first day, rats spent 60 minutes in a separate 52.5 x 26.5 x 35 cm operant box, where two levers were presented repeatedly and 45-mg food pellets were available through programmed presentations and lever responses. Two subsequent 20-minute autoshaping sessions and two 40-minute delay-reinforcement sessions took place in context A with one lever. The reinforcement program remained available during conditioning and subsequent tests. Before Study 1b retesting, the animals received renewed 60-minute instrumental practice; delayed test performance therefore reflected intervening experience and recent practice as well as elapsed time.

The program historically named VI60 implemented a response-initiated variable delay to feeder activation, not a conventional variable-interval contingency. A recorded lever input while the program awaited a response initiated selection of one of 15 programmed delays, ranging from 1.02 to 98.53 seconds, with an unweighted catalogue mean of 60.001 seconds. On completion of the delay, the controller issued a 20-millisecond feeder command and returned to waiting for a response. Additional presses during the delay were recorded but did not restart the timer. Feeder commands were not treated as direct evidence of pellet delivery or consumption. This implemented contingency defines the instrumental opportunity available during the threat procedures.

### Conditioning and subsequent tests

Study 1a conditioning began with 600 seconds in context A, followed by three 20-second, 9-kHz tones at 180-second onset-to-onset intervals. In the shock group, a 0.7-mA, one-second footshock began at tone second 19 and coterminated with the tone. Controls received the tones without shock. The recorded observation period ended at 1,140 seconds. Subsequent tests included 180 seconds in context A without a cue, a 9-kHz cue test in context B, generalization tests with 7-and 5-kHz tones, and a repeated 9-kHz cue test. Cue tests used a 180-second baseline and two unreinforced 20-second tones, with 180 seconds between their onsets, ending at 540 seconds. Study 1b repeated the context and 9-kHz cue tests in the same animals. Three additional cue acquisitions followed recording problems; they were retained as separate exposures and descriptive checks, not substituted for original acquisitions without a matching source record.

Study 2 used the same instrumental training and chamber contexts. Both groups received three white-noise and footshock pairings in context A, following a 600-second baseline. The intended shock intensity and duration were 0.7 mA and one second. On the following day, extinction animals received one unreinforced retrieval cue in context B and an extinction session scheduled approximately one hour later. No-extinction animals received corresponding context exposures without cues. Both groups subsequently received unreinforced retention tests in context B and a renewal test in context A. Three further extinction sessions in context B preceded the final retention test. During these further sessions, cues were presented to the extinction group only; the no-extinction group continued to receive context exposure without cues.

The executed schedules differed between the two Study 2 acquisition cohorts, comprising three extinction plus three no-extinction animals and six extinction plus five no-extinction animals. First-extinction sessions in the extinction group contained ten cues and lasted 1,980 seconds in the earlier cohort, but contained 18 cues and lasted 3,420 seconds in the later cohort. The initial retention and renewal tests used two cues in 540 seconds in the earlier cohort and four cues in 720 seconds in the later cohort. All three additional extinction sessions lasted 1,980 seconds, with ten cues for extinction animals. Final retention tests contained four cues in both cohorts, with different onset schedules and durations of 900 and 720 seconds. Cue times, group treatment, and analysis windows are specified by cohort in Table S1. Learning-day cohort was retained in the Study 2 statistical comparisons rather than treating these schedules as identical.

### Behavioral recording and vocal curation

FreezeFrame 5 (RRID:SCR_014429) controlled the conditioning procedure and acquired video at approximately 15 frames per second. Freezing was quantified from video-derived activity using an operator-selected motion threshold at the first inflection of the activity distribution. The analyzed historical exports recorded a minimum-bout setting of 0.25 seconds and provided freezing percentages in five-second intervals. This recorded setting, rather than the longer verbal definition in the original protocol, specifies the scores used here. Original selected exports were retained; alternative scoring versions were kept separate. The five-second percentages do not identify the exact placement of immobility within each interval.

Two Pettersson M500-384 USB ultrasonic microphones, mounted above the chambers, recorded audio with Audacity at 384 kHz. DeepSqueak detections were manually curated (40). The curation protocol used two trained assessors, with disagreements referred to a third expert. Candidate 22-kHz calls were evaluated in the 18-35-kHz band; non-call traces were excluded and missed calls were added when supported by the spectrogram. The 50-kHz criteria used 35-80 kHz and durations below 100 milliseconds. Acoustic subtype labels were not consistently retained, so pooled band counts were not assigned subtype-specific meanings. The analyses used retained call annotations, curator-designated versions and documented investigator summaries, with disagreements represented explicitly rather than resolved by the model’s fit.

GraphicState 4 controlled the instrumental program on a separate computer. Physical lever responses were defined by recorded On1A4 input onsets, including multiple responses within a five-second interval and responses during a programmed delay. Entry into the controller state named “rat press” marked response eligibility rather than a physical response (Table S1E). Repeated-onset or incomplete On/Off sequences were retained with flags rather than automatically deleted. Fecal boli were counted after each acquisition and analyzed at that level; they never formed a five-second emission channel. Audio, video and controller recordings required session-specific temporal registration, as described below.

### Behavioral observations and analysis

Behavior was analyzed at two complementary levels. Single-channel comparisons used all available measurements in the relevant experimental windows, without requiring simultaneous availability of the other channels. Temporal modeling used windows with aligned observations of freezing, 22-kHz calls, 50-kHz calls, and physical lever presses. The animal was the unit of group comparison and predictive evaluation; five-second windows and repeated recordings were observations within animals. Studies 1a and 1b involved the same 12 rats, and Study 2 contributed a separate 17 rats.

Freezing means were computed from the investigator-selected FreezeFrame exports, weighted by the recorded seconds within each analysis window. The retained exports specified a 0.25-second minimum freezing bout; alternative retained scoring settings were analyzed separately where available. Physical lever presses were counted from GraphicState input events in half-open time intervals, retaining multiple presses within a window. Entering a program state that waited for a press was not counted as a physical press. Press rates used the actual recorded duration; primary window comparisons required at least 95% coverage. Unrecorded measurements remained missing, whereas an observed interval with no event retained a count of zero.

Study 2 task summaries measured 22-kHz call duration in seconds and annotated 50-kHz event counts; these quantities retained their distinct units. Positive aggregate summaries were not distributed among five-second windows. Documented zero summaries could support zero counts only within their specified task windows. One observation policy prioritized such summaries where they disagreed with retained calls; the other gave retained event annotations priority. The reference policies used the same rows, freezing observations, physical-input counts and animal folds. An event-annotation-only analysis excluded summary-zero substitution and a separately reported recording-level zero, which was otherwise restricted to its identified recording. Source-attribution sensitivities for aggregate comparisons are specified in Table S6.

### Timing and observation coverage

Separate acquisition systems were aligned using the investigator’s processing offsets, experimental notes, GraphicState start or end markers, and identifiable sounds or saved cue annotations for each recording. These are recording-specific timing estimates, not hardware synchronization measurements. The reference lever alignment used the manual-end estimate, with the first start-switch origin examined separately. For the 32 cue-free context exposures in Study 2, operator recollection indicated that audio started a few seconds before FreezeFrame. A three-second lead defined a conditional reference, not a measured offset attributed to each session. All 32 exposures remained eligible under the same source and channel-coverage requirements as other contexts.

Joint temporal modeling included 45,300 five-second windows from 29 animals in 208 contiguous acquisition sequences. Each window required a selected freezing score, supported counts for both vocal channels, and complete physical-input recording coverage on its specified clock (Table S1C,D). Unsupported measurements remained missing, while their available channels could contribute to separate comparisons. Eligibility was determined from measurement support, not the inferred configuration. The event-annotation-only sensitivity comprised 29,130 windows from 25 animals in 117 acquisitions; it did not redefine the reference sample.

Sequence boundaries were imposed at acquisition boundaries and gaps, so transitions never joined separate recordings or unobserved intervals. To assess cue-free clock uncertainty, ten illustrative audio leads spanning 0-15 seconds and 100 deterministic draws assigning separate leads to each recording were examined. The draw range was a stress test, not a calibrated probability distribution or confidence interval. The selected parameters and preprocessing were held fixed during projection, with coverage and configuration expression reported separately. Alternative-source and feeder-linked models were also fitted independently, rather than assuming that projection alone captures refitting uncertainty.

### Group and longitudinal comparisons

Study 1 conditioning used a 0-600-second baseline and a 600-1140-second task window. Context tests used 0-180 seconds; cue and generalization tests used a 0-180-second baseline and a 180-540-second task window, including inter-cue periods. Historical investigator conditioning summaries used 700-1140 seconds and were retained separately. Original acquisitions were used for the principal comparisons; three repeated delayed-cue acquisitions were descriptive checks rather than replacements. Between-group differences in means used exhaustive animal-label permutation, conditional on exchangeability. Delayed comparisons paired each animal with its original context, first-cue, or repeated-cue measurement. Within-group changes used exhaustive paired sign flips, conditional on symmetric changes, and between-group differences in change used label permutation. Holm adjustments were applied separately to the 28 group, 20 paired-change, and 10 between-group-change comparisons within each clock version (41). Identifier-based block permutations provided a limited clustering sensitivity; the blocks were not treated as independently verified cage assignments.

Study 2 comparisons used session-specific baseline and task windows. Thirty-six group contrasts covered task levels and task-minus-baseline changes for freezing and pressing across nine phases. Exact permutations preserved the numbers of extinction and no-extinction animals within the learning-day cohorts. Sixteen paired comparisons examined the last versus first 180 seconds of each of four extinction sessions, separately by group and channel. Holm correction was applied separately to these two comparison families. Investigator USV summaries were tested by exhaustive between-group permutation, with Benjamini-Hochberg adjustment across the 18 Study 2 vocal comparisons (42); Study 1 investigator-workbook contrasts formed a separate family. These tests were conditional exchangeability analyses, not evidence that random allocation had been documented for every comparison. A post hoc sensitivity restricted these 18 vocal permutations within learning-day cohorts, preserving group counts; a separate Benjamini-Hochberg family retained all 18 comparisons. Effect estimates retain their original units: freezing percentage points, physical inputs per minute, call-duration seconds or annotated event counts. Magnitude and direction are interpreted alongside individual variation and source, clock and cohort sensitivities. Neither multiplicity-adjusted tests nor descriptive subgroup differences are treated as tests of equivalence or as independently validated cohort interactions.

Exploratory follow-up analyses included only extinction animals with temporally ordered paired acquisitions. Late-task levels and early-to-late changes in freezing or pressing were related to subsequent freezing or pressing for three transitions: first extinction to the first retention test, first extinction to renewal, and fourth extinction to the second retention test. Spearman correlations used exact outcome permutations within learning-day cohorts. All 24 specified comparisons remained in the Holm family; non-estimable comparisons contributed p = 1 to the adjustment but were reported as non-estimable, not as zero associations. These analyses did not constitute out-of-sample prognostic validation.

Configuration-to-outcome comparisons used the same three chronological pairs, with at least five eligible animals and 95% coverage of the antecedent window. The reference predictor was mass-posterior occupancy in the final 180 seconds of extinction; subsequent task-window freezing or pressing required its own channel coverage, not later four-channel coverage. All four configurations, three pairs and two outcomes remained in a 24-test Holm family within each source-policy and clock alternative, including any non-estimable entry. All nine extinction animals contributed to these reference comparisons; the use of full-data representations makes them exploratory association analyses, not independent prognostic validation. Sensitivities retained hard fractions, whole-task predictors, K=8 and the recording-specific acoustic alternative. Separate Holm families retained every component, three pairs and two outcomes within each model, earlier-clock, predictor-window and fraction definition. Constant measures remained in the multiplicity family as p = 1 but were reported as non-estimable, not zero associations.

A separate relative-clock sensitivity used feeder-associated acoustic events. Controller food-command sequences were divided into a fitting part and a held-out part, and acoustic correspondence was evaluated against local timing jitter. Twenty of 21 testable cue-free exposures showed held-out correspondence; 11 others had too few eligible commands and one did not meet the correspondence criterion. All recordings remained represented regardless of that test. A sound match corroborates controller-to-audio correspondence and response-contingent actuation; it does not identify every physical press, pellet delivery or consumption, or the audio-to-FreezeFrame origin.

For the 20 corroborated recordings, GraphicState and audio offsets were varied together using their relative correspondence. Discrepancy corrections of 0, 0.73 and 1.5 seconds examined sensitivity to the offset observed in 12 comparison recordings; these are not isolated hardware-latency calibrations. The median-correction, three-second-audio reference contained 45,304 windows and supplied independent K=4 refits. The same audio-offset scenarios were then projected with linked GraphicState offsets; unlinked records retained their operator-based mapping. Additional recording-specific GraphicState shifts of-15,-5, 0, 5 and 15 seconds stressed manual mappings. Neither sources nor offsets were selected by biological outcomes or favorable association tests.

Study 1 sensitivity used the retained processing-first and notebook-first audio mappings crossed with manual-end and first-switch lever origins. Configurations were compared on shared named-acquisition and nominal-window coordinates, with additional coverage counted separately (Figure S6; Table S9). These comparisons describe dependence on defensible timing alternatives; none measures synchronization accuracy. No common one-second correction was imposed across acquisitions.

### Temporal models and predictive evaluation

Five folds were assigned by unique animal, stratified by cohort and experimental group. All observations from the same animal, including both Study 1 series, remained in one fold. The same folds and rows were used for both vocal-source versions. The Gaussian representation used log(1 + count) for each count channel and raw freezing percentage, followed by standardization using training-fold means and standard deviations. Preprocessing parameters were then applied unchanged to held-out animals. The logarithm compresses the influence of unusually large counts while preserving their order; standardization puts the four channels on comparable scales during fitting. These choices define a statistical representation rather than establishing biological equivalence among channels, and the direct-count comparison tests a different representation.

The reference comparison fixed a four-component diagonal Gaussian HMM and an independent mixture with the same emissions at a variance floor of 0.1 in standardized units. The independent model constrained initial probabilities and every transition row to common state weights, removing temporal dependence. The same two families were compared at two through eight components with that floor under both source policies. Four-component fits at 0.01 supplied an additional variance-boundary sensitivity. Each candidate used ten starts per fold, at most 500 expectation-maximization iterations, and an absolute log-likelihood tolerance of 0.0001. Specifications were documented before these fits; the analyses were post hoc relative to the original experiments.

Gaussian fitting used continuous transformed observations. Training-set restart selection and held-out scoring instead integrated each count-channel distribution over the interval corresponding to its observed integer count. Freezing contributed a Gaussian density, including the Jacobian back to percentage units. Sequence probabilities were reset at every contiguous-sequence boundary. Eligible restarts were ranked by training-set count-mass score, with ties within 0.00000001 resolved by the lower seed. Numerical eligibility required finite, normalized and constraint-compliant parameters, nonempty decoded states, and a nondecreasing likelihood within floating-point tolerance; the final two increments had to be nonnegative and below 0.0001. All attempted fits and failed candidates were retained in the supplementary accounting.

Held-out log scores were divided by each animal’s number of observed windows and averaged across animals. Paired model differences used the same animal and observations. A candidate required an eligible fit in all five folds to enter the ranking. Across-animal standard errors and one-standard-error sets were descriptive because training samples overlapped. This was not independently nested model selection, and the fixed-reference floor was not a newly validated biological parameter. Full-data descriptions used 50 starts at four and eight components for both source policies and the event-annotation-only subset, plus four-component feeder-linked alternatives. All eligible starts contributed to stability summaries; biological descriptors were assigned from measured channel signatures, not from a preferred group or follow-up result.

A direct-count sensitivity retained the HMM dependence structure but used independent Poisson emissions for the three count channels and a Gaussian emission for freezing. Only freezing was standardized, using training-fold parameters. The four-component models used a freezing-variance floor of 0.1, matching the reference comparator, a Poisson-rate floor of 0.00000001, ten starts per fold and 50 full-data starts per source policy. Starts were selected by their own eligible training likelihood, and held-out comparisons used the same observations, animal folds and physical units as the Gaussian count-mass HMM. Poisson emissions provide a sensitivity representation, not an assumption that all biological count overdispersion is explained.

### Configuration expression and fecal boli

Configuration fractions were calculated within animal and acquisition before aggregation by phase and group. Posterior probabilities used forward-backward inference, and decoded paths used the Viterbi algorithm (43). Correspondence used optimal one-to-one assignment (44), implemented with SciPy’s linear assignment solver; normalized and adjusted mutual information used scikit-learn (45). Posterior-weighted and decoded fractions were kept separate, and partial coverage was not extrapolated to an entire acquisition. Pooled occupancy was not a normal or recovered target. Between-group task-configuration contrasts required 95% joint coverage and formed a 36-test Holm family per source-policy and clock alternative, covering four configurations and nine phases. Complete transition matrices at four and eight components included every state pair. Empirical transition probabilities included self-transitions in the outgoing denominator, never crossed an acquisition boundary or a gap, and were distinguished from fitted parameters. These descriptive matrices do not establish selective routing or a dedicated neural gate; selected-fit correspondence is not independent model transfer.

Fecal boli were analyzed as acquisition-level counts from all 153 Study 2 acquisitions. Negative-binomial generalized estimating equations used fixed dispersion alpha = 1, exchangeable animal clusters, and indicators for phase, group and learning cohort (46). Bias-reduced covariance estimates were primary (47). Freezing was expressed per ten percentage points. Occupancy models included three K=4 proportions per 0.1, omitting the configuration with the smallest empirical sum of vocal and lever counts, without consulting boli. Whole-acquisition analyses required 95% joint coverage; task-window sensitivities related task behavior to whole-acquisition boli totals. Separate freezing models used those same observations as well as the full available sample.

The 48 occupancy coefficients across two source policies, two clock references, two fraction definitions and two windows formed one Holm family. Fits required at least 30 observations and ten animal clusters, more than twice as many observations as design columns, a full-rank design, a varying outcome, and finite converged estimates. No unsuccessful model was simplified to obtain a favorable result. A separate exposure sensitivity added log recorded duration in minutes as an offset to the same models, with its own 48-coefficient accounting. Recorded duration need not equal complete chamber exposure or the time during which defecation occurred. Differences between significant and nonsignificant coefficients were not treated as evidence of superiority.

### Held-out outcome prediction

Additional prediction analyses were specified after earlier data and association results were known. They are post hoc internal evaluations, not preregistered or external validation. For both source policies and K=4 and K=8, we reused the eligible training-score-selected HMM from the target animal’s existing fold. That animal contributed neither HMM training observations nor scaler estimates. The same fold model supplied training and test features within each supervised split. Mean count-mass posterior fractions required 95% joint coverage and never linked acquisitions across a boundary or gap. Exact selected inputs, all predictions and executable replay accompany Source code 1.

Later-behavior evaluation left one of the nine extinction animals out of supervised fitting for each of the three chronological pairs and two raw outcomes described above. The first baseline used late freezing, late log(1 + inputs per minute), and last-minus-first changes in freezing and log input rate over the first and final 180 seconds. The second also used final-window log(1 + rate) for both call bands. Augmented models added all K posterior fractions. Ridge regression used training-standardized predictors, unpenalized intercept and cohort, penalty 1 as primary and 0.1 and 10 as fixed sensitivities. Predictions were bounded to 0-100% freezing or nonnegative input rate; unbounded losses were retained. RMSE and mean absolute error used original outcome units. Cohort-mean and same-response persistence forecasts were retained as benchmarks. All 144 specification comparisons, including all 48 primary comparisons, contained all nine animals. Fixed-prediction leave-one-animal loss-influence ranges were descriptive, not confidence intervals.

Physiological prediction used the existing five animal folds and matched samples of 98 whole acquisitions or 148 task windows from 17 animals. Baseline GEE models included whole-acquisition freezing, phase, group and cohort; the second raw baseline added log(1 + rate per minute) for each count channel in the specified joint window. Augmented models added K-1 posterior fractions. Continuous predictors were standardized within supervised training only. The negative-binomial family, fixed dispersion, cluster covariance and numerical eligibility rules matched the association analysis. A log recording-duration offset was primary and omission of the offset a sensitivity. Held-out negative-binomial log scores were averaged within animal and then equally over animals; absolute and squared errors were also retained. All 320 fits converged with finite estimates and no recorded warnings, yielding 32 complete matched comparisons. Specifications were not selected by their outcome scores and are not independent replications.

### Software

Analyses used Python 3.13.9, NumPy 2.3.5, SciPy 1.16.3, pandas 2.3.3, scikit-learn 1.7.2, hmmlearn 0.3.3, and statsmodels 0.14.5. Gaussian model fitting used hmmlearn with explicit variance-floor and convergence checks. The mixed Poisson-count and Gaussian-freezing model used hmmlearn inference with channel-specific emission updates. Serialization checks recomputed likelihoods, posterior probabilities and decoded paths from saved parameters; count-emission probabilities were also checked against SciPy implementations.

## Author contributions

B.Y.: Conceptualization, Methodology, Formal Analysis, Visualization, Supervision, Project Administration, Resources, Writing - Original Draft, Writing - Review & Editing. J.R.: Investigation, Methodology, Data Curation, Formal Analysis, Writing - Original Draft. X.T.W.: Conceptualization, Writing - Review & Editing.

## Data, code, and materials availability

Numerical source data accompany the figures and tables. The reconstructed observations, analyses, software environments and source provenance are publicly available in Zenodo v3.1.0 (https://doi.org/10.5281/zenodo.24030085; all-version concept DOI: https://doi.org/10.5281/zenodo.20551230). Source code 1 supplies the additional held-out outcome analyses, their exact selected inputs and complete outputs, the descriptive instrumental and resolution summaries, and code for the revised Figure 1.

## Use of artificial intelligence tools

The theoretical framework and hypotheses, experimental design, and primary data collection were developed and conducted entirely by the authors without AI assistance. After primary data collection, AI-assisted systems, including OpenAI Codex, Edison Scientific’s Kosmos, and Anthropic’s Claude Code, were used under author direction for data-structure reconciliation, execution and verification of specified analyses, code and figure checking, and language editing and document assembly. AI-generated outputs were not treated as evidence on their own. Numerical claims were checked against versioned data, executable code, and provenance records; unsupported analyses were excluded or reported as limitations. The authors retained final authority over all scientific questions, model specifications, analytical decisions, and interpretations, reviewed the outputs, and take full responsibility for the work.

## Supporting information

Supplementary Information including Figures S1-S6 and Tables S1-S12

Source code 1: Additional outcome analyses, exact inputs and complete results

## Acknowledgments

We thank members of the Laboratory for Learning and Behavioral Sciences at the School of Psychology, Fujian Normal University, for assistance with behavioral data collection, ultrasonic vocalization annotation, and data curation.

