## Supplementary Information including Figures S1-S6 and Tables S1-S12 for "Multimodal temporal modeling of behavioral reorganization across threat learning and extinction in rats"

##### Supplementary figures

Figure S1. Paired behavior at the delayed retest depends on the earlier comparison

Figure S2. Early-to-late changes within extinction-stage sessions

Figure S3. Investigator-scored vocal activity across all Study 2 phases

Figure S4. Measured signatures at eight Gaussian components

Figure S5. Measured signatures under direct-count emissions

Figure S6. Configuration sensitivity to recording-specific clocks

##### Supplementary tables

Table S1. Experimental schedules, observation reconstruction and controller measures

Table S2. Between-group behavior across initial and delayed Study 1 tests

Table S3. Within-animal changes at delayed testing in Study 1

Table S4. Extinction and no-extinction group comparisons across Study 2

Table S5. Within-session change across the four extinction-stage sessions

Table S6. Investigator-scored vocal output during Study 2

Table S7. Complete configuration transition matrices

Table S8. Post-extinction behavior and later responding within animals

Table S9. Post-extinction configurations and subsequent behavior

Table S10. Behavioral and configuration associations with fecal boli

Table S11. Held-out prediction of later behavior

Table S12. Added prediction of fecal-boli counts

**Figure S1. Paired behavior at the delayed retest depends on the earlier comparison**

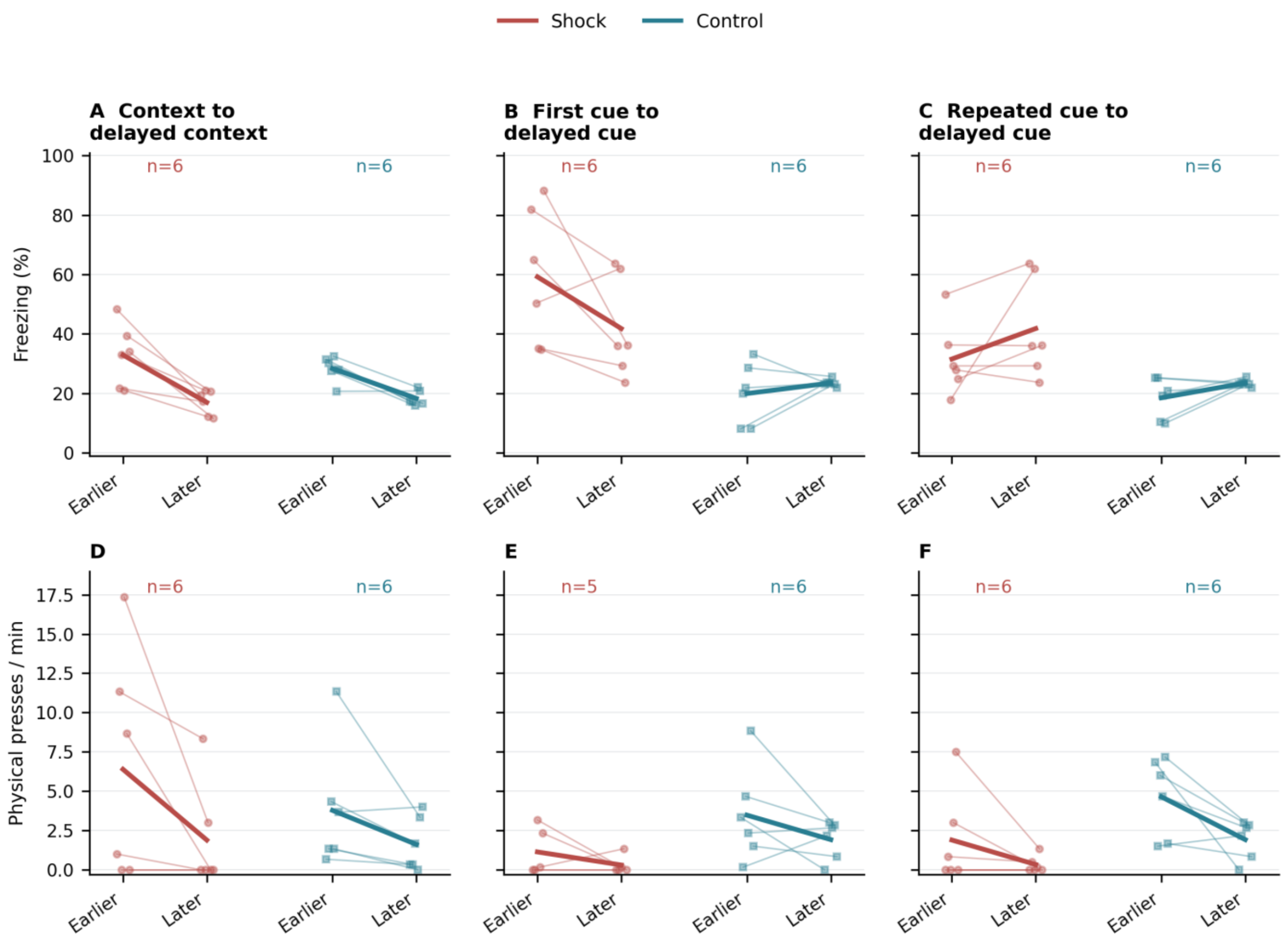


Freezing (A-C) and press-rate (D-F) changes from the original context test to delayed context (A, D), the first 9-kHz cue test to delayed cue (B, E), and the repeated 9-kHz cue test to delayed cue (C, F). Thin lines connect the same animal; thick lines connect group means. Sample sizes are shown in each panel. The first-cue press comparison contains only the five shock animals with both measurements. The same delayed-cue observations appear in two comparisons and are not independent replicates. Original acquisitions and the manual-end lever-clock estimate are used throughout. Axes share a common scale within each row. None of the 20 paired-change comparisons met the specified Holm-corrected threshold. Lower freezing at a later test is not, by definition, renewed instrumental function.

#### Figure S2. Early-to-late changes within extinction-stage sessions

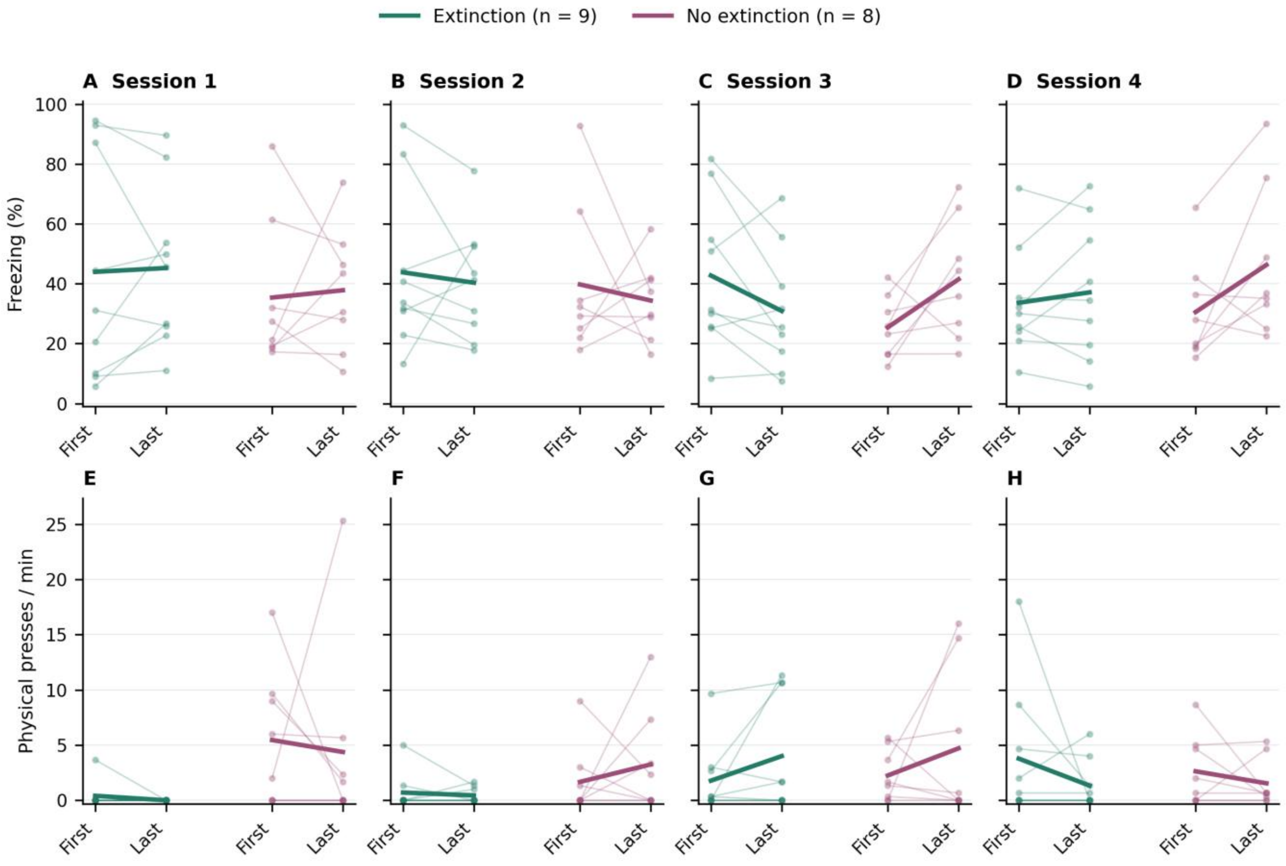


Freezing (A-D) and physical lever-press rates (E-H) in the first and last 180 seconds of each protocol-defined task window. Columns show sessions 1-4. Green lines denote the nine extinction animals and magenta lines the eight no-extinction animals receiving matched context exposure. Thin lines connect individual observations; thick lines connect group means. The comparison uses the actual start and end of each task window, including the longer first session in the later acquisition cohort, rather than imposing one common session duration. Axes share a scale within each row and retain all observed values. These within-session changes differ from comparing whole-session means across days. None of the 16 paired early-to-late tests met the Holm-corrected threshold.

**Figure S3. Investigator-scored vocal activity across all Study 2 phases**

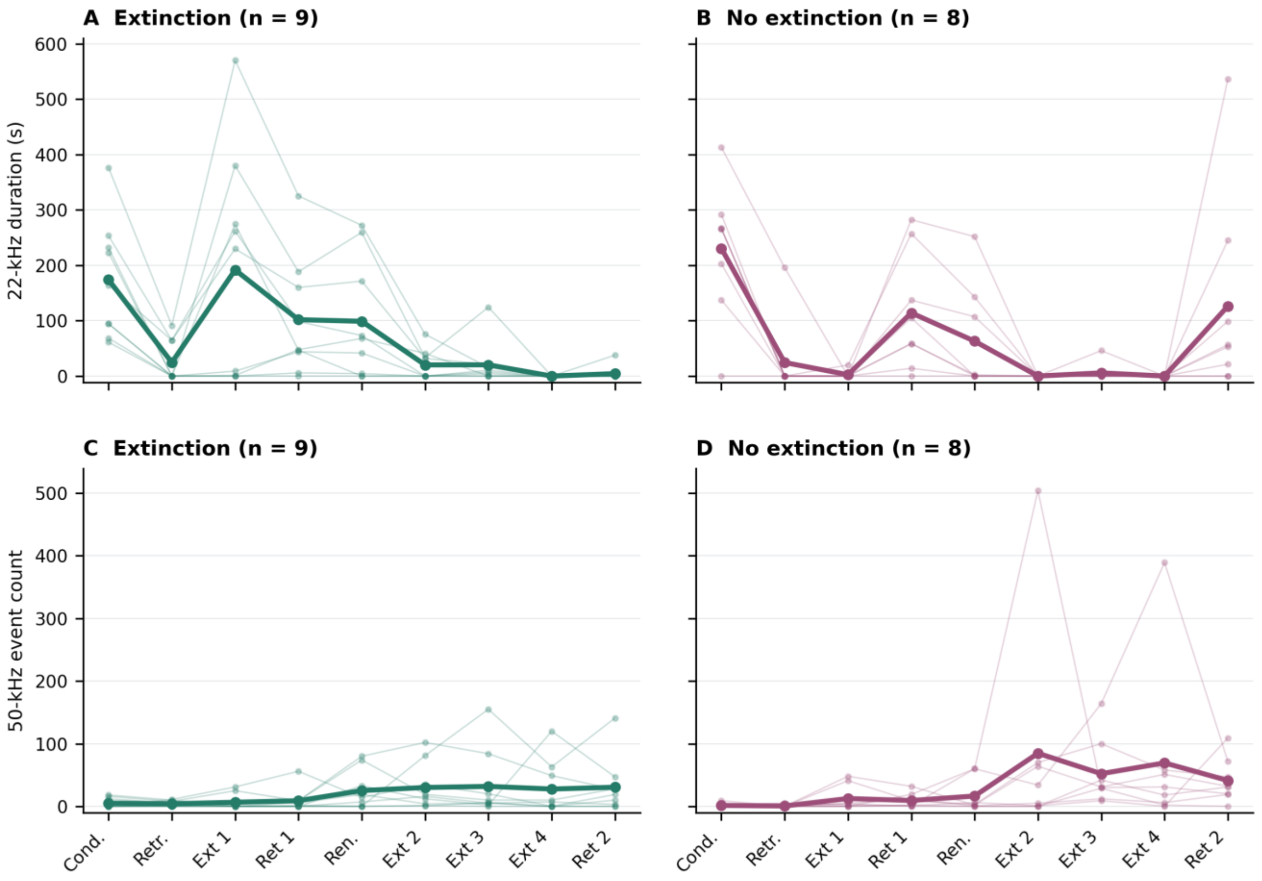


Original task-window 22-kHz durations (A, B) and annotated 50-kHz event counts (C, D). Thin lines connect values from the same animal; thick lines connect group means. All nine extinction and eight no-extinction animals contribute in every phase, regardless of eligibility for joint temporal modeling. Phase abbreviations follow Figure 3. These are totals, not rates: task-window durations differ by phase and acquisition cohort, so connecting successive phases does not establish a change in output per unit exposure. The two vocal measures also have different units and are not interchangeable. Explicit investigator-recorded zeros are retained. None of the 18 planned vocal group comparisons met the Benjamini-Hochberg false-discovery-rate threshold.

#### Figure S4. Measured signatures at eight Gaussian components

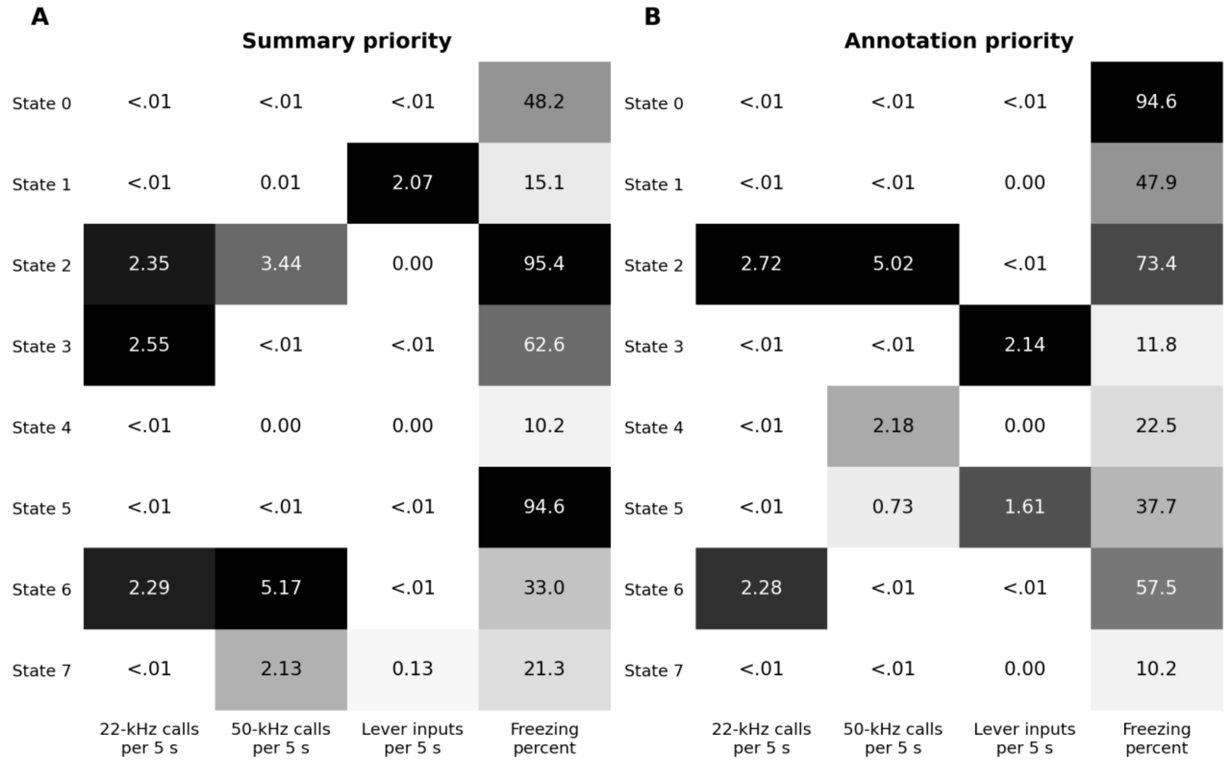


Empirical channel means in the selected K=8 Gaussian HMMs under summary priority (A) and annotation priority (B). Both use 45,300 five-second windows, the same animals, reference clocks and variance floor 0.1. Means are calculated among count-mass decoded windows, not transformed Gaussian locations. Cell shading is scaled within channel; numerical values retain their own units. State numbers belong to the individual fit and are not a common sequence of affective stages. In particular, similar low output can accompany widely different freezing levels. The eight-component fit predicts better within the tested range, but does not strictly nest into the selected four-component partition or establish eight natural states. Full restart diagnostics, stability distributions and held-out candidate scores accompany the figure. The same fit-specific indices and measured-signature names appear on the eight-component transition axes in Table S7.

#### Figure S5. Measured signatures under direct-count emissions

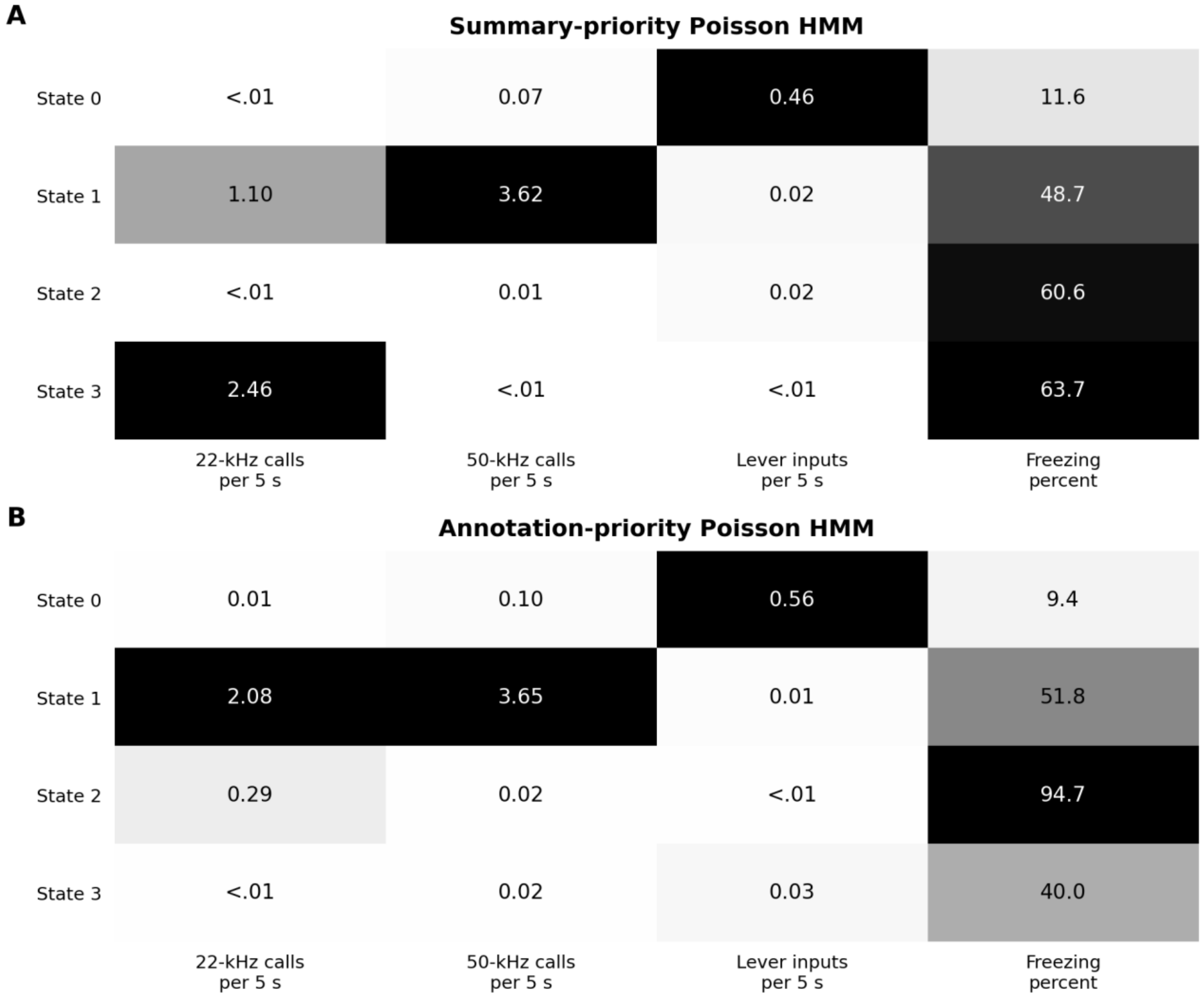


Empirical means among decoded windows of the selected Poisson-count and Gaussian-freezing HMMs, with summary priority (A) and annotation priority (B). Each uses four components, all 45,300 reference windows, a freezing-variance floor of 0.1, and the same full-data preprocessing as its cross-validation specification. The plotted values are measured observations, not fitted Poisson rates. State IDs are specific to each fit. Nonzero count means below 0.01 are shown as <.01; exact values are in the source tables. Both models retain temporal dependence. The comparison therefore changes the emission representation, not whether history within a recording is represented. Poisson counts reorganize vocal and instrumental combinations and do not identify invariant affective categories. Matched held-out comparisons appear in Figure 4B. In A, raw 0 is instrumental-rich, 1 is mixed-vocal, 2 is low-output and 3 is 22-kHz-dominant; in B, 0 is instrumental-rich, 1 is mixed-vocal, 2 is 22-kHz-rich with high freezing and 3 is low-output. These descriptions apply only to the displayed Poisson fits.

**Figure S6. Configuration sensitivity to recording-specific clocks**

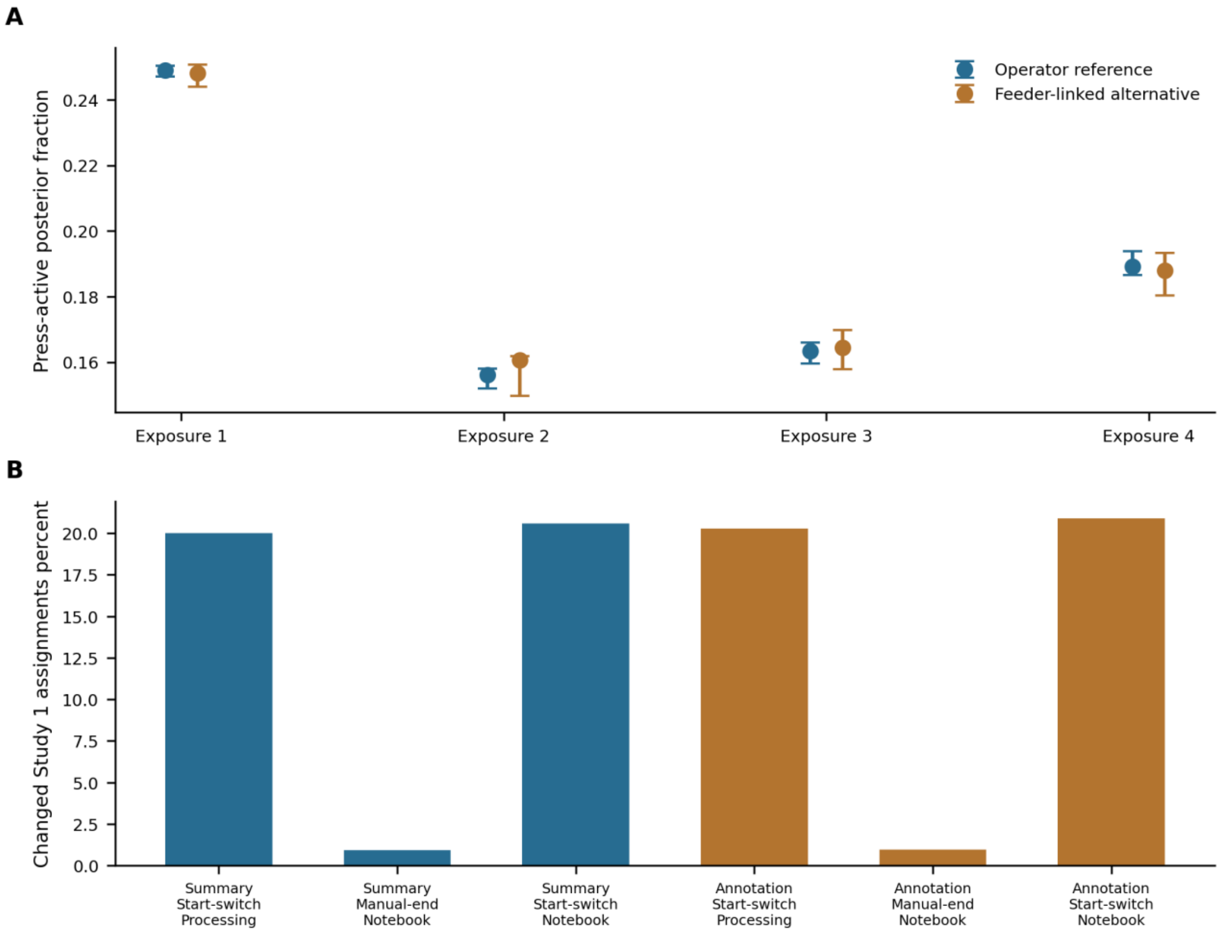


Timing alternatives are sensitivity assumptions, not measurements of synchronization error. (A) Press-active posterior occupancy in the eight cue-free context-exposure animals, averaged within each task window, under summary priority. Points show reference fractions; error bars span the declared stress realizations, not confidence intervals. The operator-reference model has 115 realizations and the feeder-linked model 335. Audio leads range from 0 to 15 seconds in common-offset checks and 100 separate per-recording draws; manual GraphicState offsets are additionally stressed at five values. In feeder-linked records, audio and controller clocks change together, with three discrepancy corrections. All 32 context-exposure recordings remain represented. (B) Fraction of assignments changed in Study 1 on the same 5,878 nominal windows when the lever origin, audio origin, or both are changed, holding the selected K=4 model fixed. Processing and notebook identify audio-source policies; start-switch and manual-end identify controller origins. The two source priorities are shown separately. These projections quantify dependence on clock assumptions, not independent confirmation of one clock or complete refitting uncertainty.

### Table S1. Experimental schedules, observation reconstruction and controller measures

Times are seconds from the start of the FreezeFrame acquisition. The nominal cue times below are rounded to the programmed second; the accompanying session-level table preserves the recorded precision. Task windows include inter-cue periods, not only cue presentation. Baseline runs from zero to the task onset, except in cue-free context tests. A no-cue group has the same analysis window as its corresponding cue group.

#### Table S1A. Studies 1a and 1b

| Phase | Cue treatment | Analysis window |
|---|---|---|
| Study 1a conditioning | Three 20-s, 9-kHz tones at 600, 780 and 960 s. Shock group: one-second shock starting at tone second 19; controls: no shock. | Baseline: 0-600 s<br>Task: 600-1140 s |
| Study 1a context | Context A without a cue. | Context: 0-180 s |
| Study 1a cue tests | Two 20-s tones at 180 and 360 s. First and repeated cue: 9 kHz; generalization tests: 7 or 5 kHz, in context B. | Baseline: 0-180 s<br>Task: 180-540 s |
| Study 1b delayed tests | The same animals received a context test and a 9-kHz cue test approximately five months later, after renewed instrumental practice. | Context: 0-180 s<br>Cue baseline: 0-180 s<br>Cue task: 180-540 s |

Each Study 1 group contained six animals. Three repeated Study 1b cue acquisitions were separate exposures and descriptive checks. Historical investigator conditioning summaries used 700-1140 s and were retained as a separate window, rather than substituted for the complete 600-1140-s task period.

##### Table S1B. Study 2

EXT denotes extinction and NOEXT denotes no extinction. Cohorts are defined by conditioning day. They are acquisition cohorts within Study 2, not independent replications of the complete experiment. Cue duration was 20 s. Only conditioning included shock; subsequent cues were unreinforced.

| Phase | Earlier cohort<br>2024-12-30<br>3 EXT and 3 NOEXT | Later cohort<br>2025-01-06<br>6 EXT and 5 NOEXT |
|---|---|---|
| Conditioning | Both groups: 3 cues<br>Onsets: 600, 780, 960 s<br>Task: 600-1140 s | Both groups: 3 cues<br>Onsets: 600, 780, 960 s<br>Task: 600-1140 s |
| Retrieval | EXT: 1 cue; NOEXT: none<br>Onsets: 180 s<br>Task: 180-360 s | EXT: 1 cue; NOEXT: none<br>Onsets: 180 s<br>Task: 180-360 s |
| Extinction 1 | EXT: 10 cues; NOEXT: none<br>Onsets: 180 to 1800, every 180 s<br>Task: 180-1980 s | EXT: 18 cues; NOEXT: none<br>Onsets: 180 to 3240, every 180 s<br>Task: 180-3420 s |
| Retention 1 | Both groups: 2 cues<br>Onsets: 180, 360 s<br>Task: 180-540 s | Both groups: 4 cues<br>Onsets: 240, 310, 480, 580 s<br>Task: 240-720 s |
| Renewal | Both groups: 2 cues<br>Onsets: 180, 360 s<br>Task: 180-540 s | Both groups: 4 cues<br>Onsets: 240, 310, 480, 580 s<br>Task: 240-720 s |
| Extinction 2 | EXT: 10 cues; NOEXT: none<br>Onsets: 180 to 1800, every 180 s<br>Task: 180-1980 s | EXT: 10 cues; NOEXT: none<br>Onsets: 180 to 1800, every 180 s<br>Task: 180-1980 s |
| Extinction 3 | EXT: 10 cues; NOEXT: none<br>Onsets: 180 to 1800, every 180 s<br>Task: 180-1980 s | EXT: 10 cues; NOEXT: none<br>Onsets: 180 to 1800, every 180 s<br>Task: 180-1980 s |
| Extinction 4 | EXT: 10 cues; NOEXT: none<br>Onsets: 180 to 1800, every 180 s<br>Task: 180-1980 s | EXT: 10 cues; NOEXT: none<br>Onsets: 180 to 1800, every 180 s<br>Task: 180-1980 s |
| Retention 2 | Both groups: 4 cues<br>Onsets: 180, 360, 540, 720 s<br>Task: 180-900 s | Both groups: 4 cues<br>Onsets: 240, 310, 480, 580 s<br>Task: 240-720 s |

First- and last-window extinction comparisons used the first and last 180 s of each stated task window. Group comparisons preserved learning-day cohort. The session-level source table includes all 153 Study 2 acquisitions, the native cue times and source identities, and the windows used in the behavioral analyses.

**Table S1C. Reconciliation of the public and revised observation matrices**

The same 29 animals contribute to both matrices. Correspondence uses named acquisitions and nominal FreezeFrame-relative intervals, not an assertion that earlier clocks or measurements were correct. The first Reviewed Preprint had 213 animal-session keys and 47,004 windows; the current reference has 208 original acquisitions and 45,300 windows. Every earlier and current row has a separate source-table entry. Study 1b reuses Study 1a animals. The first Reviewed Preprint is doi:10.7554/eLife.112910.1; its earlier analysis archive is doi:10.5281/zenodo.21883766, not the archive of the present reconstructed analysis. The present reconstruction is publicly available in Zenodo version 3.1.0 (https://doi.org/10.5281/zenodo.24030085).

| Study | Public v1 | Shared nominal windows | v1 only | Current only | Current reference |
|---|---|---|---|---|---|
| 1a | 6,948 | 5,201 | 1,747 | 107 | 5,308 |
| 1b | 1,728 | 570 | 1,158 | 0 | 570 |
| 2 | 38,328 | 35,869 | 2,459 | 3,553 | 39,422 |
| Total | 47,004 | 41,640 | 5,364 | 3,660 | 45,300 |

The row-level identity is 47,004 - 5,364 + 3,660 = 45,300: 41,640 named-acquisition and nominal-window coordinates are shared, 5,364 earlier coordinates do not meet the current source/coverage requirements, and 3,660 current coordinates were absent from the earlier complete-case matrix. These are reconstructed observations from existing recordings, not new animals or newly collected experiments. The reference retains every one of the 32 cue-free context-exposure acquisitions under explicitly approximate, recording-specific timing sensitivities. The feeder-linked median-correction alternative contains 45,304 windows, with 41,644 shared earlier coordinates; it uses a different clock specification and is reported separately as a sensitivity analysis.

Among shared nominal coordinates, 2,860 22-kHz counts, 4,458 50-kHz counts, 5,400 lever values and 2,441 freezing values differ. Differences combine source selection, timing and measurement definition; they are not isolated causal effects of individual corrections. Every changed freezing value is in Study 1a conditioning: earlier values match freeze_Xinrat0424learn.csv, whereas current values match freeze_Xinrat0424learn0822s.xlsx and the investigator master workbook. Both exports were already in the raw archive; no new freezing scoring was performed for this comparison. Earlier public artifacts remain preserved.

**Table S1D. Reference support unavailable in earlier windows**

| Unavailable reference observation | Earlier windows |
|---|---|
| Controller coverage only | 83 |
| 22-kHz value | 1,486 |
| 22-kHz value and controller coverage | 2 |
| Both vocal values | 1,344 |
| 50-kHz value | 2,437 |
| 50-kHz value and controller coverage | 12 |
| Total | 5,364 |

Availability patterns are mutually exclusive and concern the reference analysis, not whether an original recording or investigator summary was false. A missing channel value may reflect unresolved recording identity, an unavailable final event annotation, or a window outside recorded coverage. Positive aggregates are not placed artificially into five-second bins. Documented zero summaries are accepted only within their task windows, with retained-annotation and event-annotation-only sensitivities kept distinct. Of 5,364 earlier coordinates not retained, 5,281 lack a supported vocal value; 14 of those also lack full controller coverage. The other 83 lack controller coverage alone. No excluded earlier coordinate lacks a selected freezing value in this correspondence.

**Table S1E. Physical responses and controller waiting-state entries**

| Measure on the same 45,300 reference windows | Count |
|---|---|
| Registered On1A4 inputs | 10,163 |
| Entries into rat press waiting state | 1,062 |
| Windows with different event counts | 5,549 |
| Waiting entry but no input in the window | 790 |
| Physical input but no waiting entry | 4,638 |

Both event definitions were recomputed from the original controller streams on the same corrected 45,300-window clock; these are not totals from the earlier 47,004-window matrix. All 208 contributing acquisitions contain the response-initiated delay flow described in Methods. Whole-recording counts in the source tables additionally include intervals outside the joint sample. An entry into the state named rat press arms or rearms the schedule; an On1A4 input registers a lever response. The two events are not interchangeable.

A waiting-state entry marks the program becoming ready to accept the next delay-initiating response, not the animal starting to wait. Its occurrence and rate depend on previous responses, the programmed delay, and schedule-cycle completion. It can therefore carry information about the task sequence while failing to measure current response vigor: presses made during the delay are recorded as inputs but do not create new waiting-state entries. A response-triggered exit can identify a delay-initiating press; the preceding entry is not that press. Neither variable establishes reward expectation, pellet consumption or goal-directed control. Treating the old entry variable as press count could favor schedule-related rather than action-defined configurations, but this operational distinction alone does not identify the cause of every change in the fitted partition.

#### Table S2. Between-group behavior across initial and delayed Study 1 tests

The same 12 animals contributed the original and approximately five-month delayed observations. Means compare six shock with six control animals except the first-cue press record (five shock animals). Freezing is in percent and pressing in physical inputs per minute. Task-minus-baseline differences retain these units. Differences are shock minus control. Exact two-sided animal-label permutation p values and Holm-adjusted p values are reported for the 28-test family within each timing version. Diff. is an unstandardized effect estimate, not an index of statistical significance. Its magnitude is read together with the individual observations and source-table standard deviations; failure to cross an adjusted threshold does not establish a negligible effect.

##### Table S2A. Task-window or whole-context behavior

| Phase | Measure | n S/C | Shock mean | Control mean | Diff. | p | Holm p |
|---|---|---|---|---|---|---|---|
| Conditioning | Freezing | 6/6 | 28.87 | 22.54 | 6.33 | 0.3983 | 1.0000 |
| Conditioning | Pressing | 6/6 | 0.80 | 3.81 | -3.02 | 0.0065 | 0.1558 |
| Context | Freezing | 6/6 | 32.94 | 28.40 | 4.54 | 0.3550 | 1.0000 |
| Context | Pressing | 6/6 | 6.39 | 3.78 | 2.61 | 0.4545 | 1.0000 |
| First 9-kHz cue | Freezing | 6/6 | 59.21 | 19.97 | 39.24 | 0.0022 | 0.0606 |
| First 9-kHz cue | Pressing | 5/6 | 1.13 | 3.47 | -2.34 | 0.1342 | 1.0000 |
| 7-kHz cue | Freezing | 6/6 | 43.80 | 21.63 | 22.17 | 0.0260 | 0.5455 |
| 7-kHz cue | Pressing | 6/6 | 1.08 | 2.47 | -1.39 | 0.3723 | 1.0000 |
| 5-kHz cue | Freezing | 6/6 | 29.99 | 20.72 | 9.27 | 0.1385 | 1.0000 |
| 5-kHz cue | Pressing | 6/6 | 1.50 | 3.22 | -1.72 | 0.3247 | 1.0000 |
| Repeated 9-kHz cue | Freezing | 6/6 | 31.59 | 18.53 | 13.05 | 0.0368 | 0.6623 |
| Repeated 9-kHz cue | Pressing | 6/6 | 1.89 | 4.64 | -2.75 | 0.1212 | 1.0000 |
| Delayed context | Freezing | 6/6 | 17.01 | 18.32 | -1.32 | 0.4978 | 1.0000 |
| Delayed context | Pressing | 6/6 | 1.89 | 1.61 | 0.28 | 0.9242 | 1.0000 |
| Delayed 9-kHz cue | Freezing | 6/6 | 41.82 | 23.53 | 18.29 | 0.0043 | 0.1126 |
| Delayed 9-kHz cue | Pressing | 6/6 | 0.33 | 1.92 | -1.58 | 0.0281 | 0.5628 |

##### Table S2B. Changes from the within-acquisition baseline

| Phase | Measure | n S/C | Shock mean | Control mean | Diff. | p | Holm p |
|---|---|---|---|---|---|---|---|
| Conditioning | Freezing | 6/6 | 3.15 | -2.65 | 5.80 | 0.6342 | 1.0000 |
| Conditioning | Pressing | 6/6 | -8.10 | -0.70 | -7.40 | 0.0108 | 0.2489 |
| First 9-kHz cue | Freezing | 6/6 | 42.97 | 1.37 | 41.60 | 0.0022 | 0.0606 |
| First 9-kHz cue | Pressing | 5/6 | -1.20 | 1.03 | -2.23 | 0.3009 | 1.0000 |
| 7-kHz cue | Freezing | 6/6 | 22.78 | 3.29 | 19.49 | 0.0043 | 0.1126 |
| 7-kHz cue | Pressing | 6/6 | -0.97 | -1.36 | 0.39 | 0.7922 | 1.0000 |

| Phase | Measure | n S/C | Shock mean | Control mean | Diff. | p | Holm p |
|---|---|---|---|---|---|---|---|
| 5-kHz cue | Freezing | 6/6 | 4.18 | -2.61 | 6.79 | 0.1623 | 1.0000 |
| 5-kHz cue | Pressing | 6/6 | -1.00 | -1.06 | 0.06 | 0.9913 | 1.0000 |
| Repeated 9-kHz cue | Freezing | 6/6 | 5.21 | -1.39 | 6.60 | 0.0325 | 0.6169 |
| Repeated 9-kHz cue | Pressing | 6/6 | -0.39 | 1.42 | -1.81 | 0.2403 | 1.0000 |
| Delayed 9-kHz cue | Freezing | 6/6 | 20.77 | 5.83 | 14.94 | 0.0152 | 0.3333 |
| Delayed 9-kHz cue | Pressing | 6/6 | -1.72 | -0.81 | -0.92 | 0.5476 | 1.0000 |

##### Table S2C. Pressing under the alternative start-switch clock origin

| Phase | Endpoint | n S/C | Shock mean | Control mean | Diff. | p | Holm p |
|---|---|---|---|---|---|---|---|
| Conditioning | Task - baseline | 6/6 | -7.55 | 0.42 | -7.98 | 0.0043 | 0.1126 |
| Conditioning | Task | 6/6 | 1.13 | 4.91 | -3.78 | 0.0087 | 0.1991 |
| Context | Task | 6/6 | 6.22 | 3.61 | 2.61 | 0.4459 | 1.0000 |
| First 9-kHz cue | Task - baseline | 5/6 | -0.90 | 1.25 | -2.15 | 0.2316 | 1.0000 |
| First 9-kHz cue | Task | 5/6 | 1.23 | 3.53 | -2.29 | 0.1775 | 1.0000 |
| 7-kHz cue | Task - baseline | 6/6 | -1.11 | -1.50 | 0.39 | 0.8268 | 1.0000 |
| 7-kHz cue | Task | 6/6 | 1.06 | 2.39 | -1.33 | 0.3853 | 1.0000 |
| 5-kHz cue | Task - baseline | 6/6 | -1.03 | -1.08 | 0.06 | 0.9957 | 1.0000 |
| 5-kHz cue | Task | 6/6 | 1.36 | 3.19 | -1.83 | 0.2749 | 1.0000 |
| Repeated 9-kHz cue | Task - baseline | 6/6 | -0.47 | 1.33 | -1.81 | 0.3074 | 1.0000 |
| Repeated 9-kHz cue | Task | 6/6 | 1.81 | 4.33 | -2.53 | 0.1190 | 1.0000 |
| Delayed context | Task | 6/6 | 1.28 | 1.33 | -0.06 | 1.0000 | 1.0000 |
| Delayed 9-kHz cue | Task - baseline | 6/6 | -1.31 | -0.81 | -0.50 | 0.6818 | 1.0000 |
| Delayed 9-kHz cue | Task | 6/6 | 0.47 | 1.92 | -1.44 | 0.0346 | 0.6580 |

A and B use the session-specific manual-end estimate. C changes only the lever-clock origin; freezing is unchanged. Both timing versions include all 28 tests in their separate Holm families. Source tables additionally retain standard deviations, identifier-block permutation sensitivities and exact permutation counts. Identifier blocks are not independently verified cage assignments.

##### Table S2D. Investigator-scored 50-kHz vocal activity during conditioning

Original workbook event summaries compare the shock and no-shock groups in the baseline and conditioning periods. These are totals in different windows, not exposure-standardized rates. Differences are shock minus control. Exact two-sided animal-label permutations and Benjamini-Hochberg q values use the complete 49-comparison Study 1 workbook family, which includes other phases and measures. The group comparison does not by itself distinguish conditioning from habituation or establish exploration.

| Period | n S/C | Shock mean | Control mean | Diff. | p | BH q |
|---|---|---|---|---|---|---|
| Baseline | 6/6 | 64.17 | 73.67 | -9.50 | 0.7641 | 0.9600 |
| Conditioning | 6/6 | 10.17 | 10.67 | -0.50 | 0.9892 | 1.0000 |

##### Table S2E. Configuration occupancy during Study 1a conditioning

Study 1a original conditioning acquisitions, six shocked and six control animals. Baseline is 0-600 s and task is 600-1140 s on the reference FreezeFrame clock. Cells are animal-averaged fractions from the selected summary-priority K=4 Gaussian model and count-mass decoding. Decoded values count assigned windows; posterior values average component probabilities. The descriptions identify measured combinations, not subjective states. These data are included in model fitting, so the comparison is descriptive, not independent transfer. Individual values, both source priorities, four and eight components, and all retained Study 1 clock alternatives are in the source data.

| Configuration | Group | n | Baseline decoded % | Task decoded % | Baseline posterior % | Task posterior % |
|---|---|---|---|---|---|---|
| Low output lower freezing | Shock | 6 | 40.1 | 33.2 | 37.3 | 31.6 |
| Low output lower freezing | Control | 6 | 43.8 | 62.8 | 41.3 | 59.0 |
| Low output higher freezing | Shock | 6 | 14.2 | 9.6 | 14.7 | 10.2 |
| Low output higher freezing | Control | 6 | 11.5 | 13.6 | 11.9 | 14.6 |
| Vocal active | Shock | 6 | 18.2 | 52.9 | 18.1 | 53.0 |
| Vocal active | Control | 6 | 22.2 | 3.2 | 22.1 | 4.0 |
| Press active | Shock | 6 | 27.5 | 4.3 | 29.9 | 5.3 |
| Press active | Control | 6 | 22.5 | 20.4 | 24.7 | 22.4 |

### Table S3. Within-animal changes at delayed testing in Study 1

Each animal was paired with its own earlier measurement. The delayed cue test is compared separately with the first and repeated 9-kHz tests; these comparisons reuse the delayed observation and are not independent replications. Changes are delayed minus earlier. Freezing means are percentages and their changes are percentage points; pressing is in physical inputs per minute. A uses the manual-end lever-clock estimate. Exact paired sign-flip tests form a 20-comparison Holm family.

#### Table S3A. Paired changes within each experimental group

| Earlier test / endpoint | Measure / group | n | Earlier mean | Delayed mean | Change | p | Holm p |
|---|---|---|---|---|---|---|---|
| Context<br>Task | Freezing<br>Control | 6 | 28.40 | 18.32 | -10.08 | 0.0625 | 1.0000 |
| Context<br>Task | Freezing<br>Shock | 6 | 32.94 | 17.01 | -15.94 | 0.0312 | 0.6250 |
| Context<br>Task | Pressing<br>Control | 6 | 3.78 | 1.61 | -2.17 | 0.0625 | 1.0000 |
| Context<br>Task | Pressing<br>Shock | 6 | 6.39 | 1.89 | -4.50 | 0.1250 | 1.0000 |
| First cue<br>Task - baseline | Freezing<br>Control | 6 | 1.37 | 5.83 | 4.46 | 0.0938 | 1.0000 |
| First cue<br>Task - baseline | Freezing<br>Shock | 6 | 42.97 | 20.77 | -22.20 | 0.0938 | 1.0000 |
| First cue<br>Task - baseline | Pressing<br>Control | 6 | 1.03 | -0.81 | -1.83 | 0.5938 | 1.0000 |
| First cue<br>Task - baseline | Pressing<br>Shock | 5 | -1.20 | -1.83 | -0.63 | 0.5000 | 1.0000 |
| First cue<br>Task | Freezing<br>Control | 6 | 19.97 | 23.53 | 3.56 | 0.3438 | 1.0000 |
| First cue<br>Task | Freezing<br>Shock | 6 | 59.21 | 41.82 | -17.39 | 0.1250 | 1.0000 |
| First cue<br>Task | Pressing<br>Control | 6 | 3.47 | 1.92 | -1.56 | 0.2812 | 1.0000 |
| First cue<br>Task | Pressing<br>Shock | 5 | 1.13 | 0.30 | -0.83 | 0.5000 | 1.0000 |
| Repeated cue<br>Task - baseline | Freezing<br>Control | 6 | -1.39 | 5.83 | 7.21 | 0.0312 | 0.6250 |
| Repeated cue<br>Task - baseline | Freezing<br>Shock | 6 | 5.21 | 20.77 | 15.55 | 0.0625 | 1.0000 |
| Repeated cue<br>Task - baseline | Pressing<br>Control | 6 | 1.42 | -0.81 | -2.22 | 0.3125 | 1.0000 |
| Repeated cue<br>Task - baseline | Pressing<br>Shock | 6 | -0.39 | -1.72 | -1.33 | 0.5000 | 1.0000 |
| Repeated cue<br>Task | Freezing<br>Control | 6 | 18.53 | 23.53 | 5.00 | 0.2188 | 1.0000 |

| Earlier test / endpoint | Measure / group | n | Earlier mean | Delayed mean | Change | p | Holm p |
|---|---|---|---|---|---|---|---|
| Repeated cue Task | Freezing Shock | 6 | 31.59 | 41.82 | 10.24 | 0.2188 | 1.0000 |
| Repeated cue Task | Pressing Control | 6 | 4.64 | 1.92 | -2.72 | 0.0625 | 1.0000 |
| Repeated cue Task | Pressing Shock | 6 | 1.89 | 0.33 | -1.56 | 0.2500 | 1.0000 |

#### Table S3B. Between-group differences in paired change

| Earlier test / endpoint | Measure | n S/C | Shock change | Control change | Diff. | p | Holm p |
|---|---|---|---|---|---|---|---|
| Context Task | Freezing | 6/6 | -15.94 | -10.08 | -5.86 | 0.2078 | 1.0000 |
| Context Task | Pressing | 6/6 | -4.50 | -2.17 | -2.33 | 0.4286 | 1.0000 |
| First cue Task - baseline | Freezing | 6/6 | -22.20 | 4.46 | -26.66 | 0.0173 | 0.1732 |
| First cue Task - baseline | Pressing | 5/6 | -0.63 | -1.83 | 1.20 | 0.7879 | 1.0000 |
| First cue Task | Freezing | 6/6 | -17.39 | 3.56 | -20.95 | 0.0563 | 0.5065 |
| First cue Task | Pressing | 5/6 | -0.83 | -1.56 | 0.72 | 0.6299 | 1.0000 |
| Repeated cue Task - baseline | Freezing | 6/6 | 15.55 | 7.21 | 8.34 | 0.0801 | 0.6407 |
| Repeated cue Task - baseline | Pressing | 6/6 | -1.33 | -2.22 | 0.89 | 0.7035 | 1.0000 |
| Repeated cue Task | Freezing | 6/6 | 10.24 | 5.00 | 5.24 | 0.6537 | 1.0000 |
| Repeated cue Task | Pressing | 6/6 | -1.56 | -2.72 | 1.17 | 0.4545 | 1.0000 |

B compares shock minus control changes by exact animal-label permutation, with Holm correction across a separate 10-comparison family.

##### Table S3C. Paired press changes under the alternative start-switch origin

| Earlier test / endpoint | Group | n | Earlier mean | Delayed mean | Change | p | Holm p |
|---|---|---|---|---|---|---|---|
| Context<br>Task | Control | 6 | 3.61 | 1.33 | -2.28 | 0.0312 | 0.6250 |
| Context<br>Task | Shock | 6 | 6.22 | 1.28 | -4.94 | 0.1250 | 1.0000 |
| First cue<br>Task - baseline | Control | 6 | 1.25 | -0.81 | -2.06 | 0.5312 | 1.0000 |
| First cue<br>Task - baseline | Shock | 5 | -0.90 | -1.43 | -0.53 | 0.5625 | 1.0000 |
| First cue<br>Task | Control | 6 | 3.53 | 1.92 | -1.61 | 0.2500 | 1.0000 |
| First cue<br>Task | Shock | 5 | 1.23 | 0.43 | -0.80 | 0.5000 | 1.0000 |
| Repeated cue<br>Task - baseline | Control | 6 | 1.33 | -0.81 | -2.14 | 0.3125 | 1.0000 |
| Repeated cue<br>Task - baseline | Shock | 6 | -0.47 | -1.31 | -0.83 | 0.7500 | 1.0000 |
| Repeated cue<br>Task | Control | 6 | 4.33 | 1.92 | -2.42 | 0.0625 | 1.0000 |
| Repeated cue<br>Task | Shock | 6 | 1.81 | 0.47 | -1.33 | 0.3750 | 1.0000 |

##### Table S3D. Between-group press-change sensitivity

| Earlier test / endpoint | n S/C | Shock change | Control change | Diff. | p | Holm p |
|---|---|---|---|---|---|---|
| Context<br>Task | 6/6 | -4.94 | -2.28 | -2.67 | 0.4048 | 1.0000 |
| First cue<br>Task - baseline | 5/6 | -0.53 | -2.06 | 1.52 | 0.7229 | 1.0000 |
| First cue<br>Task | 5/6 | -0.80 | -1.61 | 0.81 | 0.6212 | 1.0000 |
| Repeated cue<br>Task - baseline | 6/6 | -0.83 | -2.14 | 1.31 | 0.5476 | 1.0000 |
| Repeated cue<br>Task | 6/6 | -1.33 | -2.42 | 1.08 | 0.4740 | 1.0000 |

C and D retain the original 20- and 10-test correction families, respectively, including unchanged freezing comparisons. Original acquisitions are used throughout; repeated acquisitions are retained separately in the source data. None of these comparisons establishes a causal effect of elapsed time.

### Table S4. Extinction and no-extinction group comparisons across Study 2

All nine extinction and eight no-extinction animals contribute to these channel-specific comparisons. Freezing is in percent and physical pressing in inputs per minute. Differences are extinction minus no extinction. Task windows follow each executed schedule (Table S1); task-minus-baseline changes are computed within acquisitions. Exact two-sided label permutations preserve group counts within learning-day cohorts. Holm correction includes all 36 comparisons. Diff. reports an unstandardized effect estimate in percentage points or physical inputs per minute. The groups already differed during conditioning before extinction, and cohort-specific schedules differ; these contrasts are not isolated causal estimates of extinction efficacy. Individual measurements and learning-day identifiers accompany the full comparison families.

#### Table S4A. Freezing

| Phase | Endpoint | Extinction mean | Context-exposure mean | Diff. | p | Holm p |
|---|---|---|---|---|---|---|
| Conditioning | Task | 35.921 | 57.410 | -21.489 | 0.0646 | 1.0000 |
| Conditioning | Task minus baseline | 23.753 | 43.714 | -19.961 | 0.1160 | 1.0000 |
| Retrieval | Task | 59.780 | 46.056 | 13.725 | 0.3982 | 1.0000 |
| Retrieval | Task minus baseline | 41.536 | 16.637 | 24.899 | 0.0773 | 1.0000 |
| Extinction 1 | Task | 48.830 | 33.818 | 15.012 | 0.1569 | 1.0000 |
| Extinction 1 | Task minus baseline | 29.404 | 13.219 | 16.185 | 0.0847 | 1.0000 |
| Retention 1 | Task | 63.977 | 73.039 | -9.062 | 0.3865 | 1.0000 |
| Retention 1 | Task minus baseline | 45.058 | 51.132 | -6.074 | 0.4987 | 1.0000 |
| Renewal | Task | 46.818 | 67.534 | -20.716 | 0.1174 | 1.0000 |
| Renewal | Task minus baseline | 29.084 | 39.839 | -10.755 | 0.3739 | 1.0000 |
| Extinction 2 | Task | 38.744 | 34.303 | 4.441 | 0.5192 | 1.0000 |
| Extinction 2 | Task minus baseline | 18.117 | 1.844 | 16.273 | 0.0235 | 0.7515 |
| Extinction 3 | Task | 33.089 | 33.319 | -0.230 | 0.9731 | 1.0000 |
| Extinction 3 | Task minus baseline | 13.260 | 11.051 | 2.209 | 0.6642 | 1.0000 |
| Extinction 4 | Task | 32.545 | 35.664 | -3.118 | 0.7130 | 1.0000 |
| Extinction 4 | Task minus baseline | 11.270 | 9.522 | 1.748 | 0.7929 | 1.0000 |
| Retention 2 | Task | 31.371 | 67.768 | -36.397 | 0.0065 | 0.2208 |
| Retention 2 | Task minus baseline | 12.772 | 43.042 | -30.270 | 0.0220 | 0.7250 |

#### Table S4B. Physical pressing

| Phase | Endpoint | Extinction mean | Context-exposure mean | Diff. | p | Holm p |
|---|---|---|---|---|---|---|
| Conditioning | Task | 0.074 | 0.097 | -0.023 | 0.7009 | 1.0000 |
| Conditioning | Task minus baseline | -9.548 | -8.353 | -1.195 | 0.6636 | 1.0000 |
| Retrieval | Task | 0.000 | 1.417 | -1.417 | 0.2273 | 1.0000 |
| Retrieval | Task minus baseline | -1.556 | 1.000 | -2.556 | 0.0606 | 1.0000 |

| Phase | Endpoint | Extinction mean | Context-exposure mean | Diff. | p | Holm p |
|---|---|---|---|---|---|---|
| Extinction 1 | Task | 0.245 | 5.667 | -5.422 | 0.0055 | 0.1932 |
| Extinction 1 | Task minus baseline | -1.607 | 3.792 | -5.399 | 0.0037 | 0.1325 |
| Retention 1 | Task | 0.028 | 0.016 | 0.012 | 1.0000 | 1.0000 |
| Retention 1 | Task minus baseline | -1.556 | -1.496 | -0.059 | 0.9610 | 1.0000 |
| Renewal | Task | 0.194 | 0.000 | 0.194 | 1.0000 | 1.0000 |
| Renewal | Task minus baseline | -6.574 | -1.073 | -5.501 | 0.0555 | 1.0000 |
| Extinction 2 | Task | 1.381 | 2.533 | -1.152 | 0.2002 | 1.0000 |
| Extinction 2 | Task minus baseline | -1.285 | 2.117 | -3.402 | 0.0240 | 0.7515 |
| Extinction 3 | Task | 2.385 | 3.433 | -1.048 | 0.4871 | 1.0000 |
| Extinction 3 | Task minus baseline | -0.985 | 1.642 | -2.627 | 0.1158 | 1.0000 |
| Extinction 4 | Task | 3.193 | 4.463 | -1.270 | 0.6307 | 1.0000 |
| Extinction 4 | Task minus baseline | -1.622 | 2.838 | -4.460 | 0.1069 | 1.0000 |
| Retention 2 | Task | 2.653 | 0.000 | 2.653 | 0.0303 | 0.9091 |
| Retention 2 | Task minus baseline | -3.838 | -1.240 | -2.598 | 0.4460 | 1.0000 |

#### Table S4C. Alternative retained retrieval-freezing score

| Endpoint | Extinction mean | No-extinction mean | Diff. | p | Holm p |
|---|---|---|---|---|---|
| Task | 59.96 | 46.06 | 13.90 | 0.3924 | 1.0000 |
| Task - baseline | 41.67 | 16.64 | 25.04 | 0.0769 | 1.0000 |

C substitutes the alternative retained score for one extinction animal at retrieval. The other 34 group contrasts are numerically unchanged, but the Holm family is recalculated across all 36 tests. This is a scoring sensitivity, not a score selected for its statistical result. Full-precision results and individual observations accompany the tables.

### Table S5. Within-session change across the four extinction-stage sessions

First and last refer to the first and last 180 seconds of each animal's protocol-defined task window. The extinction group received cue presentations; the no-extinction group received matched context exposure without cues. Changes are last minus first. Freezing is in percent and physical pressing in inputs per minute. Exact paired sign-flip tests assume symmetric changes; all 16 comparisons form one Holm family.

#### Early and late task behavior

| Session | Group and measure | n | First mean | Last mean | Change | p | Holm p |
|---|---|---|---|---|---|---|---|
| Extinction 1 | EXT freezing | 9 | 43.964 | 45.231 | 1.267 | 0.8633 | 1.0000 |
| Extinction 1 | EXT pressing | 9 | 0.407 | 0.000 | -0.407 | 1.0000 | 1.0000 |
| Extinction 1 | NOEXT freezing | 8 | 35.373 | 37.805 | 2.432 | 0.8281 | 1.0000 |
| Extinction 1 | NOEXT pressing | 8 | 5.458 | 4.375 | -1.083 | 0.6875 | 1.0000 |
| Extinction 2 | EXT freezing | 9 | 43.788 | 40.335 | -3.453 | 0.6055 | 1.0000 |
| Extinction 2 | EXT pressing | 9 | 0.704 | 0.445 | -0.259 | 0.7500 | 1.0000 |
| Extinction 2 | NOEXT freezing | 8 | 39.752 | 34.393 | -5.359 | 0.6641 | 1.0000 |
| Extinction 2 | NOEXT pressing | 8 | 1.667 | 3.250 | 1.583 | 0.5312 | 1.0000 |
| Extinction 3 | EXT freezing | 9 | 42.734 | 30.901 | -11.833 | 0.1016 | 1.0000 |
| Extinction 3 | EXT pressing | 9 | 1.778 | 4.000 | 2.222 | 0.2188 | 1.0000 |
| Extinction 3 | NOEXT freezing | 8 | 25.430 | 41.478 | 16.048 | 0.0703 | 1.0000 |
| Extinction 3 | NOEXT pressing | 8 | 2.250 | 4.709 | 2.459 | 0.4531 | 1.0000 |
| Extinction 4 | EXT freezing | 9 | 33.624 | 37.113 | 3.490 | 0.4336 | 1.0000 |
| Extinction 4 | EXT pressing | 9 | 3.778 | 1.334 | -2.444 | 0.3750 | 1.0000 |
| Extinction 4 | NOEXT freezing | 8 | 30.550 | 46.272 | 15.722 | 0.1094 | 1.0000 |
| Extinction 4 | NOEXT pressing | 8 | 2.625 | 1.542 | -1.083 | 0.5000 | 1.0000 |

Animals, not five-second bins, are the paired units. These contrasts concern within-session change and do not substitute for between-session change. Zero values are observed zeros, not missing recordings.

### Table S6. Investigator-scored vocal output during Study 2

Original task-window summaries contain all nine extinction and eight no-extinction animals in every phase. The 22-kHz measure is total call duration in seconds; the 50-kHz measure is annotated event count. Totals are not exposure-standardized rates, and window duration differs across phases and acquisition cohorts. Differences are extinction minus no extinction. Exact two-sided animal-label permutations are reported with Benjamini-Hochberg q values across the 18 vocal comparisons.

#### Task-window vocal group contrasts

| Phase | Vocal measure | Extinction mean | No-extinction mean | Diff. | p | BH q |
|---|---|---|---|---|---|---|
| Conditioning | 22-kHz duration | 174.22 | 230.67 | -56.46 | 0.3193 | 0.7638 |
| Conditioning | 50-kHz count | 5.22 | 1.50 | 3.72 | 0.2344 | 0.7638 |
| Retrieval | 22-kHz duration | 24.50 | 24.47 | 0.02 | 1.0000 | 1.0000 |
| Retrieval | 50-kHz count | 4.11 | 0.62 | 3.49 | 0.0627 | 0.3706 |
| Extinction 1 | 22-kHz duration | 191.84 | 2.49 | 189.35 | 0.0173 | 0.1559 |
| Extinction 1 | 50-kHz count | 6.56 | 12.50 | -5.94 | 0.4760 | 0.7638 |
| Retention 1 | 22-kHz duration | 101.85 | 114.16 | -12.30 | 0.8148 | 0.9778 |
| Retention 1 | 50-kHz count | 9.00 | 9.38 | -0.38 | 0.9660 | 1.0000 |
| Renewal | 22-kHz duration | 98.96 | 63.02 | 35.94 | 0.4807 | 0.7638 |
| Renewal | 50-kHz count | 25.22 | 16.50 | 8.72 | 0.5516 | 0.7638 |
| Extinction 2 | 22-kHz duration | 20.17 | 0.04 | 20.13 | 0.0824 | 0.3706 |
| Extinction 2 | 50-kHz count | 30.11 | 84.75 | -54.64 | 0.5304 | 0.7638 |
| Extinction 3 | 22-kHz duration | 20.32 | 5.75 | 14.57 | 0.4596 | 0.7638 |
| Extinction 3 | 50-kHz count | 31.89 | 52.12 | -20.24 | 0.4482 | 0.7638 |
| Extinction 4 | 22-kHz duration | 0.00 | 0.00 | 0.00 | 1.0000 | 1.0000 |
| Extinction 4 | 50-kHz count | 27.67 | 69.38 | -41.71 | 0.5158 | 0.7638 |
| Retention 2 | 22-kHz duration | 4.45 | 126.37 | -121.92 | 0.0041 | 0.0740 |
| Retention 2 | 50-kHz count | 30.56 | 41.25 | -10.69 | 0.5976 | 0.7683 |

Recorded zeros apply only within supported task windows; positive aggregates remain available for these comparisons even without resolved event attribution. Two 22-kHz entries have uncertain attribution: learning EXT2R, 94.54587 s, and first retention EXT2B, 47.72452 s. Their event files reproduce the reported totals but internally identify EXT2G.wav and EXT2R.wav; original-waveform checks support those identities. Neither entry is reassigned or changed to zero. Omitting only these two cells preserves the complete 18-test, cohort-restricted Benjamini-Hochberg family. Minimum q is 0.0662 with either treatment. These tests establish neither absent vocal contrasts nor resolution of the historical attribution.

### Table S7. Complete configuration transition matrices

Each cell gives fitted transition probability above empirical adjacent-transition probability from count-mass decoding. N out is the number of observed outgoing adjacent pairs, including persistence. Only contiguous observations within an acquisition are paired. Raw component numbers belong to each fit and need not match between source policies or resolutions. All counts, both decoding conventions, selected signatures and group-, phase- and study-resolved transitions are retained in source data. These descriptions do not establish selective fear-goal routing. F denotes freezing. Every axis pairs the original raw index with its measured signature; identical numbers in different fits do not indicate matching configurations.

##### Measured signatures and indices at four components

| Source priority | ID | Observed description | 22-kHz / 5 s | 50-kHz / 5 s | Inputs / 5 s | Freezing % |
|---|---|---|---|---|---|---|
| Summary | 0 | Low output lower freezing | 0.000 | 0.000 | 0.000 | 15.8 |
| Summary | 1 | Vocal active | 1.610 | 1.703 | 0.004 | 52.9 |
| Summary | 2 | Low output higher freezing | 0.001 | 0.002 | 0.003 | 74.0 |
| Summary | 3 | Press active | 0.001 | 0.173 | 1.876 | 14.8 |
| Annotation | 0 | Low output higher freezing | 0.002 | 0.004 | 0.003 | 74.1 |
| Annotation | 1 | Press active | 0.002 | 0.130 | 1.975 | 14.9 |
| Annotation | 2 | Low output lower freezing | 0.001 | 0.014 | 0.000 | 15.8 |
| Annotation | 3 | Vocal active | 1.650 | 2.547 | 0.004 | 52.5 |

##### 4 components with investigator-summary priority

| From and to | 0<br>Low output lower F | 1<br>Vocal active | 2<br>Low output higher F | 3<br>Press active | N out |
|---|---|---|---|---|---|
| 0<br>Low output lower F | 0.766<br>0.784 | 0.032<br>0.021 | 0.091<br>0.080 | 0.111<br>0.114 | 20,682 |
| 1<br>Vocal active | 0.089<br>0.057 | 0.824<br>0.896 | 0.062<br>0.027 | 0.025<br>0.020 | 6,821 |
| 2<br>Low output higher F | 0.138<br>0.125 | 0.030<br>0.014 | 0.810<br>0.841 | 0.023<br>0.020 | 12,218 |
| 3<br>Press active | 0.452<br>0.468 | 0.018<br>0.015 | 0.032<br>0.028 | 0.497<br>0.489 | 5,371 |

##### 4 components with retained-annotation priority

| From and to | 0<br>Low output higher F | 1<br>Press active | 2<br>Low output lower F | 3<br>Vocal active | N out |
|---|---|---|---|---|---|
| 0<br>Low output higher F | 0.807<br>0.837 | 0.023<br>0.019 | 0.138<br>0.126 | 0.033<br>0.018 | 12,323 |
| 1<br>Press active | 0.032<br>0.026 | 0.501<br>0.517 | 0.448<br>0.441 | 0.018<br>0.015 | 5,099 |
| 2<br>Low output lower F | 0.091<br>0.081 | 0.111<br>0.100 | 0.766<br>0.798 | 0.032<br>0.021 | 20,985 |
| 3<br>Vocal active | 0.067<br>0.034 | 0.025<br>0.020 | 0.088<br>0.057 | 0.819<br>0.889 | 6,685 |

#### Table S7 continued at 8 components

##### 8 components with investigator-summary priority

| From and to | 0 Low output 48% F | 1 Press active 15% F | 2 Mixed vocal 95% F | 3 22-kHz 63% F | 4 Low output 10% F | 5 Low output 95% F | 6 Mixed vocal 33% F | 7 50-kHz 21% F | N out |
|---|---|---|---|---|---|---|---|---|---|
| 0 Low output 48% F | 0.710 0.761 | 0.042 0.037 | 0.001 0.001 | 0.004 0.003 | 0.157 0.128 | 0.058 0.050 | 0.001 0.001 | 0.026 0.018 | 12,375 |
| 1 Press active 15% F | 0.103 0.081 | 0.498 0.515 | 0.000 0.000 | 0.000 0.000 | 0.368 0.380 | 0.001 0.001 | 0.000 0.000 | 0.030 0.023 | 4,681 |
| 2 Mixed vocal 95% F | 0.015 0.009 | 0.000 0.000 | 0.809 0.820 | 0.000 0.000 | 0.003 0.000 | 0.144 0.140 | 0.028 0.030 | 0.002 0.001 | 896 |
| 3 22-kHz 63% F | 0.018 0.011 | 0.001 0.000 | 0.000 0.000 | 0.922 0.947 | 0.024 0.018 | 0.035 0.023 | 0.000 0.000 | 0.000 0.000 | 2,978 |
| 4 Low output 10% F | 0.145 0.115 | 0.104 0.105 | 0.000 0.000 | 0.006 0.005 | 0.696 0.740 | 0.001 0.000 | 0.004 0.002 | 0.044 0.033 | 15,338 |
| 5 Low output 95% F | 0.135 0.113 | 0.008 0.006 | 0.022 0.021 | 0.017 0.011 | 0.001 0.000 | 0.815 0.847 | 0.001 0.001 | 0.001 0.000 | 5,215 |
| 6 Mixed vocal 33% F | 0.029 0.022 | 0.002 0.002 | 0.089 0.077 | 0.000 0.000 | 0.070 0.034 | 0.001 0.000 | 0.801 0.859 | 0.009 0.005 | 555 |
| 7 50-kHz 21% F | 0.109 0.070 | 0.068 0.054 | 0.000 0.000 | 0.000 0.000 | 0.228 0.166 | 0.002 0.002 | 0.001 0.001 | 0.590 0.706 | 3,054 |

##### 8 components with retained-annotation priority

| From and to | 0 Low output 95% F | 1 Low output 48% F | 2 Mixed vocal 73% F | 3 Press active 12% F | 4 50-kHz 22% F | 5 Press + vocal 38% F | 6 22-kHz 57% F | 7 Low output 10% F | N out |
|---|---|---|---|---|---|---|---|---|---|
| 0 Low output 95% F | 0.807 0.840 | 0.135 0.110 | 0.026 0.025 | 0.001 0.000 | 0.001 0.000 | 0.009 0.010 | 0.020 0.014 | 0.001 0.000 | 5,250 |
| 1 Low output 48% F | 0.059 0.051 | 0.711 0.762 | 0.003 0.002 | 0.033 0.029 | 0.020 0.014 | 0.012 0.012 | 0.005 0.003 | 0.157 0.127 | 12,366 |
| 2 Mixed vocal 73% F | 0.064 0.062 | 0.016 0.011 | 0.889 0.904 | 0.000 0.000 | 0.005 0.004 | 0.001 0.001 | 0.002 0.000 | 0.023 0.017 | 2,270 |
| 3 Press active 12% F | 0.000 0.000 | 0.086 0.074 | 0.000 0.000 | 0.492 0.487 | 0.017 0.011 | 0.026 0.020 | 0.001 0.000 | 0.377 0.407 | 4,234 |
| 4 50-kHz 22% F | 0.002 0.002 | 0.086 0.052 | 0.005 0.003 | 0.039 0.029 | 0.630 0.722 | 0.052 0.060 | 0.001 0.000 | 0.185 0.131 | 2,722 |
| 5 Press + vocal 38% F | 0.021 0.017 | 0.215 0.206 | 0.003 0.002 | 0.229 0.197 | 0.146 0.159 | 0.162 0.189 | 0.002 0.002 | 0.223 0.230 | 661 |
| 6 22-kHz 57% F | 0.053 0.038 | 0.032 0.016 | 0.004 0.003 | 0.001 0.000 | 0.001 0.000 | 0.000 0.000 | 0.877 0.920 | 0.032 0.022 | 2,127 |
| 7 Low output 10% F | 0.001 0.000 | 0.145 0.113 | 0.005 0.004 | 0.102 0.104 | 0.036 0.024 | 0.006 0.005 | 0.006 0.004 | 0.700 0.746 | 15,462 |

#### Table S8. Post-extinction behavior and later responding within animals

Exploratory Spearman associations use the nine extinction animals with temporally ordered paired acquisitions. Late level is the final 180 s of the extinction task; change is last minus first 180 s. Later outcomes are task freezing or registered physical-input rate. Exact permutations preserve acquisition cohort. All 24 comparisons remain in one Holm family, including non-estimable measures. These are within-sample associations, not independently validated predictions. The raw extinction-animal outcomes are unchanged by the new feeder-link alternative, which concerns cue-free control exposures.

##### Table S8A. First extinction to first retention

| Earlier measure | Earlier endpoint | Later measure | n | rho | p | Holm p |
|---|---|---|---|---|---|---|
| Freezing | Late level | Freezing | 9 | 0.417 | 0.2454 | 1.0000 |
| Freezing | Late level | Pressing | 9 | -0.137 | 0.8333 | 1.0000 |
| Freezing | Late - early | Freezing | 9 | -0.500 | 0.1898 | 1.0000 |
| Freezing | Late - early | Pressing | 9 | 0.411 | 0.5000 | 1.0000 |
| Pressing | Late level | Freezing | 9 | NE | NE | NE |
| Pressing | Late level | Pressing | 9 | NE | NE | NE |
| Pressing | Late - early | Freezing | 9 | 0.274 | 0.8333 | 1.0000 |
| Pressing | Late - early | Pressing | 9 | -1.000 | 0.1667 | 1.0000 |

##### Table S8B. First extinction to renewal

| Earlier measure | Earlier endpoint | Later measure | n | rho | p | Holm p |
|---|---|---|---|---|---|---|
| Freezing | Late level | Freezing | 9 | 0.150 | 0.7074 | 1.0000 |
| Freezing | Late level | Pressing | 9 | -0.137 | 0.8333 | 1.0000 |
| Freezing | Late - early | Freezing | 9 | -0.633 | 0.0840 | 1.0000 |
| Freezing | Late - early | Pressing | 9 | 0.411 | 0.5000 | 1.0000 |
| Pressing | Late level | Freezing | 9 | NE | NE | NE |
| Pressing | Late level | Pressing | 9 | NE | NE | NE |
| Pressing | Late - early | Freezing | 9 | 0.411 | 0.5000 | 1.0000 |
| Pressing | Late - early | Pressing | 9 | -1.000 | 0.1667 | 1.0000 |

##### Table S8C. Fourth extinction to second retention

| Earlier measure | Earlier endpoint | Later measure | n | rho | p | Holm p |
|---|---|---|---|---|---|---|
| Freezing | Late level | Freezing | 9 | 0.833 | 0.0662 | 1.0000 |
| Freezing | Late level | Pressing | 9 | -0.731 | 0.0296 | 0.6815 |
| Freezing | Late - early | Freezing | 9 | 0.433 | 0.3551 | 1.0000 |
| Freezing | Late - early | Pressing | 9 | -0.357 | 0.3741 | 1.0000 |
| Pressing | Late level | Freezing | 9 | -0.365 | 0.3611 | 1.0000 |
| Pressing | Late level | Pressing | 9 | 0.877 | 0.0083 | 0.2000 |
| Pressing | Late - early | Freezing | 9 | 0.096 | 0.8519 | 1.0000 |
| Pressing | Late - early | Pressing | 9 | -0.200 | 0.6204 | 1.0000 |

NE denotes a non-estimable correlation because one measure was constant. Such comparisons contributed p = 1 only to multiplicity accounting; they are not reported as zero correlations or evidence of no association. Ties and sparse pressing are retained.

#### Table S9. Post-extinction configurations and subsequent behavior

Each comparison pairs mass-posterior configuration occupancy in the final 180 s of extinction with later task freezing or pressing in the same animal. At least 95% antecedent joint coverage and the later outcome's own channel coverage are required; later four-channel availability is not required. All nine extinction animals are included for each of the three chronological pairs under both source priorities and both selected clock references. No configuration association meets its 24-test Holm threshold. The press-active fourth-extinction to second-retention pressing association has rho = 0.923, exact p = 0.0028 and Holm p = 0.0667 under both source priorities; these are the same nine animals. The full-data representation and small sample preclude independent prognosis.

##### Table S9A. Paired sample support

| Selected model and clock | Earlier phase | Paired animals | Excluded animal |
|---|---|---|---|
| Summary / operator | Extinction 1 | 9 / 9 | None |
| Summary / operator | Extinction 4 | 9 / 9 | None |
| Annotation / operator | Extinction 1 | 9 / 9 | None |
| Annotation / operator | Extinction 4 | 9 / 9 | None |
| Summary / feeder | Extinction 1 | 9 / 9 | None |
| Summary / feeder | Extinction 4 | 9 / 9 | None |
| Annotation / feeder | Extinction 1 | 9 / 9 | None |
| Annotation / feeder | Extinction 4 | 9 / 9 | None |

Counts refer to paired animals, not bins or restart fits. The reference K=4 final-window posterior analysis has 96 estimable comparisons across source and clock references, each with nine animals. The complete sensitivity retains whole-task and final-window predictors, hard and posterior fractions, K=4 and K=8, and reference and recording-specific acoustic antecedent alignments. Its 1,536 comparisons form 48 separate 24- or 48-test families. Of these, 1,388 are estimable and 148 contain a constant measure; none meets its specified Holm threshold. Every slot includes nine source-supported animal pairs. Constant measures contribute p = 1 only to correction accounting and are reported as non-estimable. No missing channel is treated as zero.

Complete source tables contain all individual fractions, paired observations and sensitivity tests. Timing alternatives are conditional mappings, not independent replications. Configuration identities belong to each selected representation. The printed tables show final-window K=4 posterior associations; whole-task, hard-fraction, K=8 and acoustic alternatives retain their complete separate families in source data. The use of full-data representations precludes independent prognostic validation.

#### Table S9B. Final 180 seconds configuration fractions

Summary-priority K=4 model with mass-posterior final-window fractions. Each of four configurations is related to each later raw outcome for all three pairs. All 24 comparisons form one Holm family. Freezing is percent and pressing is registered inputs per minute. Exact Spearman permutations preserve acquisition cohort. F denotes freezing; the raw index is retained beside each empirical configuration name.

| Pair | Configuration (raw index) | Later outcome | n | rho | p | Holm p |
|---|---|---|---|---|---|---|
| Ext 1 to Ret 1 | 0 Low output, lower F | Freezing | 9 | -0.467 | 0.2062 | 1.0000 |
| Ext 1 to Ret 1 | 0 Low output, lower F | Pressing | 9 | 0.137 | 0.8333 | 1.0000 |
| Ext 1 to Ret 1 | 1 Vocal active | Freezing | 9 | -0.017 | 0.9780 | 1.0000 |
| Ext 1 to Ret 1 | 1 Vocal active | Pressing | 9 | 0.137 | 0.8333 | 1.0000 |
| Ext 1 to Ret 1 | 2 Low output, higher F | Freezing | 9 | 0.450 | 0.2056 | 1.0000 |
| Ext 1 to Ret 1 | 2 Low output, higher F | Pressing | 9 | -0.137 | 0.8333 | 1.0000 |
| Ext 1 to Ret 1 | 3 Press active | Freezing | 9 | -0.467 | 0.2062 | 1.0000 |
| Ext 1 to Ret 1 | 3 Press active | Pressing | 9 | 0.137 | 0.8333 | 1.0000 |
| Ext 1 to Renewal | 0 Low output, lower F | Freezing | 9 | -0.233 | 0.5472 | 1.0000 |
| Ext 1 to Renewal | 0 Low output, lower F | Pressing | 9 | 0.137 | 0.8333 | 1.0000 |
| Ext 1 to Renewal | 1 Vocal active | Freezing | 9 | -0.033 | 0.9456 | 1.0000 |
| Ext 1 to Renewal | 1 Vocal active | Pressing | 9 | 0.137 | 0.8333 | 1.0000 |
| Ext 1 to Renewal | 2 Low output, higher F | Freezing | 9 | 0.250 | 0.5125 | 1.0000 |
| Ext 1 to Renewal | 2 Low output, higher F | Pressing | 9 | -0.137 | 0.8333 | 1.0000 |
| Ext 1 to Renewal | 3 Press active | Freezing | 9 | -0.233 | 0.5472 | 1.0000 |
| Ext 1 to Renewal | 3 Press active | Pressing | 9 | 0.137 | 0.8333 | 1.0000 |
| Ext 4 to Ret 2 | 0 Low output, lower F | Freezing | 9 | -0.733 | 0.0690 | 1.0000 |
| Ext 4 to Ret 2 | 0 Low output, lower F | Pressing | 9 | 0.618 | 0.0954 | 1.0000 |
| Ext 4 to Ret 2 | 1 Vocal active | Freezing | 9 | -0.067 | 0.8940 | 1.0000 |
| Ext 4 to Ret 2 | 1 Vocal active | Pressing | 9 | -0.270 | 0.5056 | 1.0000 |
| Ext 4 to Ret 2 | 2 Low output, higher F | Freezing | 9 | 0.833 | 0.0662 | 1.0000 |
| Ext 4 to Ret 2 | 2 Low output, higher F | Pressing | 9 | -0.731 | 0.0296 | 0.6815 |
| Ext 4 to Ret 2 | 3 Press active | Freezing | 9 | -0.650 | 0.1449 | 1.0000 |
| Ext 4 to Ret 2 | 3 Press active | Pressing | 9 | 0.923 | 0.0028 | 0.0667 |

##### Table S9C. Retained-annotation-priority final-window associations

The same definitions and 24-test multiplicity family as Table S9B, with the retained-annotation-priority K=4 model. State numbers identify this fit rather than matching raw numbers in another model. Feeder-linked estimates are available in full-precision source data. F denotes freezing; the names, rather than the raw indices, allow the corresponding configurations to be recognized across panels.

| Pair | Configuration (raw index) | Later outcome | n | rho | p | Holm p |
|---|---|---|---|---|---|---|
| Ext 1 to Ret 1 | 0 Low output, higher F | Freezing | 9 | 0.517 | 0.1569 | 1.0000 |
| Ext 1 to Ret 1 | 0 Low output, higher F | Pressing | 9 | -0.137 | 0.8333 | 1.0000 |
| Ext 1 to Ret 1 | 1 Press active | Freezing | 9 | -0.467 | 0.2062 | 1.0000 |
| Ext 1 to Ret 1 | 1 Press active | Pressing | 9 | 0.137 | 0.8333 | 1.0000 |
| Ext 1 to Ret 1 | 2 Low output, lower F | Freezing | 9 | -0.467 | 0.2062 | 1.0000 |
| Ext 1 to Ret 1 | 2 Low output, lower F | Pressing | 9 | 0.137 | 0.8333 | 1.0000 |
| Ext 1 to Ret 1 | 3 Vocal active | Freezing | 9 | 0.050 | 0.9000 | 1.0000 |
| Ext 1 to Ret 1 | 3 Vocal active | Pressing | 9 | 0.000 | 1.0000 | 1.0000 |
| Ext 1 to Renewal | 0 Low output, higher F | Freezing | 9 | 0.317 | 0.4076 | 1.0000 |
| Ext 1 to Renewal | 0 Low output, higher F | Pressing | 9 | -0.137 | 0.8333 | 1.0000 |
| Ext 1 to Renewal | 1 Press active | Freezing | 9 | -0.233 | 0.5472 | 1.0000 |
| Ext 1 to Renewal | 1 Press active | Pressing | 9 | 0.137 | 0.8333 | 1.0000 |
| Ext 1 to Renewal | 2 Low output, lower F | Freezing | 9 | -0.233 | 0.5472 | 1.0000 |
| Ext 1 to Renewal | 2 Low output, lower F | Pressing | 9 | 0.137 | 0.8333 | 1.0000 |
| Ext 1 to Renewal | 3 Vocal active | Freezing | 9 | 0.067 | 0.8731 | 1.0000 |
| Ext 1 to Renewal | 3 Vocal active | Pressing | 9 | 0.000 | 1.0000 | 1.0000 |
| Ext 4 to Ret 2 | 0 Low output, higher F | Freezing | 9 | 0.833 | 0.0662 | 1.0000 |
| Ext 4 to Ret 2 | 0 Low output, higher F | Pressing | 9 | -0.731 | 0.0296 | 0.6815 |
| Ext 4 to Ret 2 | 1 Press active | Freezing | 9 | -0.650 | 0.1449 | 1.0000 |
| Ext 4 to Ret 2 | 1 Press active | Pressing | 9 | 0.923 | 0.0028 | 0.0667 |
| Ext 4 to Ret 2 | 2 Low output, lower F | Freezing | 9 | -0.733 | 0.0690 | 1.0000 |
| Ext 4 to Ret 2 | 2 Low output, lower F | Pressing | 9 | 0.618 | 0.0954 | 1.0000 |
| Ext 4 to Ret 2 | 3 Vocal active | Freezing | 9 | 0.117 | 0.7940 | 1.0000 |
| Ext 4 to Ret 2 | 3 Vocal active | Pressing | 9 | -0.348 | 0.3472 | 1.0000 |

**Table S9D. Cue-free context timing sensitivity**

The 32 context-exposure recordings contain no scheduled auditory CS anchor. Operator recollection supports an approximate audio lead of a few seconds, not recovered per-session numeric calibration. The reference assumes 3 s; ten common-offset checks and 100 per-recording draws span 0-15 s. A uniform draw is a sensitivity device, not a probability model for the unknown clock.

| Context-exposure phase | Recordings retained | Testable feeder sequences | Held-out matches | Audio-lead assumption |
|---|---|---|---|---|
| Extinction 1 | 8 | 5 | 4 | 3 s; stress 0-15 s |
| Extinction 2 | 8 | 4 | 4 | 3 s; stress 0-15 s |
| Extinction 3 | 8 | 5 | 5 | 3 s; stress 0-15 s |
| Extinction 4 | 8 | 7 | 7 | 3 s; stress 0-15 s |

Twenty of 21 testable recordings show held-out correspondence between controller feeder commands and sound; the 11 underpowered recordings and one unmatched recording are retained. The acoustic link constrains controller-to-audio timing, while audio-to-video timing remains uncertain. In linked records both offsets move together. The three corrections 0, 0.73 and 1.5 s reflect an empirical discrepancy sensitivity, not isolated hardware latency. Across the four fixed selected models there are 900 projected realizations, all retaining the 32 context exposures. Fraction ranges are not confidence intervals. The reference model and coupled alternative were refitted independently; projection alone does not capture every refitting uncertainty.

### Table S10. Behavioral and configuration associations with fecal boli

Negative-binomial GEE uses exchangeable animal clusters, fixed alpha = 1, and phase, group and acquisition-cohort adjustment. Coefficients are log rate ratios per ten percentage points, with bias-reduced, unadjusted 95% confidence intervals. Occupancy contrasts redistribute fraction from the lowest-count reference, holding other included proportions fixed. All 48 coefficients across source priorities, clocks, fraction definitions and windows form one Holm family; raw-freezing models are separate comparisons. All models include 17 animals. A difference in significance between estimates does not establish a difference between those estimates.

#### Table S10A. Summary priority with lower-freezing low-output reference

| Window and fraction | Measured term (raw index) | n | Coef. | 95% CI | p | Holm p |
|---|---|---|---|---|---|---|
| Whole recording<br>all raw freezing | Freezing | 153 | 0.115 | 0.005 to 0.224 | 0.0398 | NE |
| Whole recording<br>posterior fraction | 1 Vocal active | 98 | 0.013 | -0.151 to 0.178 | 0.8742 | 1.0000 |
| Whole recording<br>posterior fraction | 2 Low output, higher F | 98 | -0.060 | -0.236 to 0.116 | 0.5059 | 1.0000 |
| Whole recording<br>posterior fraction | 3 Press active | 98 | -0.314 | -0.762 to 0.134 | 0.1699 | 1.0000 |
| Whole recording<br>hard fraction | 1 Vocal active | 98 | 0.020 | -0.140 to 0.180 | 0.8069 | 1.0000 |
| Whole recording<br>hard fraction | 2 Low output, higher F | 98 | -0.054 | -0.225 to 0.116 | 0.5306 | 1.0000 |
| Whole recording<br>hard fraction | 3 Press active | 98 | -0.336 | -0.794 to 0.123 | 0.1511 | 1.0000 |
| Whole recording<br>matched freezing | Freezing | 98 | 0.083 | -0.030 to 0.196 | 0.1503 | NE |
| Task<br>posterior fraction | 1 Vocal active | 148 | 0.080 | -0.037 to 0.197 | 0.1803 | 1.0000 |
| Task<br>posterior fraction | 2 Low output, higher F | 148 | 0.026 | -0.103 to 0.155 | 0.6916 | 1.0000 |
| Task<br>posterior fraction | 3 Press active | 148 | -0.121 | -0.550 to 0.309 | 0.5821 | 1.0000 |
| Task<br>hard fraction | 1 Vocal active | 148 | 0.079 | -0.034 to 0.193 | 0.1698 | 1.0000 |
| Task<br>hard fraction | 2 Low output, higher F | 148 | 0.025 | -0.099 to 0.148 | 0.6959 | 1.0000 |
| Task<br>hard fraction | 3 Press active | 148 | -0.130 | -0.589 to 0.329 | 0.5795 | 1.0000 |
| Task support<br>matched freezing | Freezing | 148 | 0.112 | 0.004 to 0.219 | 0.0414 | NE |

At least 95% joint coverage is required. Whole-recording analyses include 98 acquisitions and task analyses 148. Task fractions are associated with whole-acquisition boli totals, not time-localized defecation. Both posterior and hard fractions remain in the 48-term family. This printed panel uses the operator-clock reference; the feeder-linked counterpart is retained in source data. Omitted reference: raw 0, low output with lower freezing (15.8%). F denotes freezing in the term labels.

#### Table S10B. Annotation priority with higher-freezing low-output reference

| Window and fraction | Measured term (raw index) | n | Coef. | 95% CI | p | Holm p |
|---|---|---|---|---|---|---|
| Whole recording posterior fraction | 1 Press active | 98 | -0.245 | -0.667 to 0.177 | 0.2548 | 1.0000 |
| Whole recording posterior fraction | 2 Low output, lower F | 98 | 0.055 | -0.120 to 0.231 | 0.5369 | 1.0000 |
| Whole recording posterior fraction | 3 Vocal active | 98 | 0.073 | -0.062 to 0.207 | 0.2893 | 1.0000 |
| Whole recording hard fraction | 1 Press active | 98 | -0.241 | -0.670 to 0.188 | 0.2706 | 1.0000 |
| Whole recording hard fraction | 2 Low output, lower F | 98 | 0.043 | -0.130 to 0.216 | 0.6274 | 1.0000 |
| Whole recording hard fraction | 3 Vocal active | 98 | 0.067 | -0.070 to 0.205 | 0.3371 | 1.0000 |
| Whole recording matched freezing | Freezing | 98 | 0.083 | -0.030 to 0.196 | 0.1503 | NE |
| Task posterior fraction | 1 Press active | 148 | -0.141 | -0.498 to 0.215 | 0.4367 | 1.0000 |
| Task posterior fraction | 2 Low output, lower F | 148 | -0.028 | -0.155 to 0.100 | 0.6722 | 1.0000 |
| Task posterior fraction | 3 Vocal active | 148 | 0.055 | -0.028 to 0.138 | 0.1957 | 1.0000 |
| Task hard fraction | 1 Press active | 148 | -0.137 | -0.523 to 0.249 | 0.4866 | 1.0000 |
| Task hard fraction | 2 Low output, lower F | 148 | -0.029 | -0.151 to 0.092 | 0.6348 | 1.0000 |
| Task hard fraction | 3 Vocal active | 148 | 0.054 | -0.029 to 0.138 | 0.2042 | 1.0000 |
| Task support matched freezing | Freezing | 148 | 0.112 | 0.004 to 0.219 | 0.0414 | NE |

Definitions and coverage are the same as Table S10A: 98 whole-recording and 148 task acquisitions, representing all 17 animals. Reference state choice is determined only by measured channel means, without consulting boli. Full source data retain both clock references and all attempted model terms. Omitted reference: raw 0, low output with higher freezing (74.1%). F denotes freezing in the term labels.

#### Table S10C. Whole-recording exposure-duration sensitivity

| Observation policy and measure | Measured term (raw index) | n | Coef. | 95% CI | p | Holm p |
|---|---|---|---|---|---|---|
| All raw / freezing | Freezing | 153 | 0.100 | -0.006 to 0.206 | 0.0643 | NE |
| Summary / posterior | 1 Vocal active | 98 | 0.033 | -0.124 to 0.190 | 0.6780 | 1.0000 |
| Summary / posterior | 2 Low output, higher F | 98 | -0.052 | -0.221 to 0.117 | 0.5453 | 1.0000 |
| Summary / posterior | 3 Press active | 98 | -0.302 | -0.767 to 0.162 | 0.2023 | 1.0000 |
| Summary / freezing | Freezing | 98 | 0.079 | -0.027 to 0.185 | 0.1422 | NE |
| Annotation / posterior | 1 Press active | 98 | -0.241 | -0.675 to 0.193 | 0.2768 | 1.0000 |
| Annotation / posterior | 2 Low output, lower F | 98 | 0.047 | -0.121 to 0.215 | 0.5820 | 1.0000 |
| Annotation / posterior | 3 Vocal active | 98 | 0.084 | -0.043 to 0.212 | 0.1939 | 1.0000 |

The exposure sensitivity adds log recorded duration in minutes to the same models. Duration may differ from total time in the chamber and does not identify the actual time of defecation. Its 48 configuration coefficients form a separate Holm family; none meets the corrected threshold. The printed entries use whole-recording posterior fractions, with corresponding freezing estimates. References follow the smallest empirical sum of count-channel means, not the same freezing category across fits: the operator-clock reference is raw state 0 in both source priorities, lower-freezing under summary priority and higher-freezing under annotation priority. Each coefficient is therefore interpreted relative to its own model reference, not as an identical cross-model biological contrast. The complete source tables retain task, hard-fraction and feeder-linked exposure models rather than selecting estimates by p value. Each configuration term carries its empirical name and raw index; F denotes freezing.

#### Table S11. Held-out prediction of later behavior

Each comparison contains all nine extinction animals. A target animal is excluded from HMM/scaler and supervised training. Raw baseline F/P uses late freezing and pressing and their early-to-late changes; four-channel adds both call rates. Ridge penalty 1 is primary. RMSE is in freezing percentage points or physical inputs per minute. Persistence carries forward the same late-extinction response. All 144 comparisons, including fixed penalties 0.1 and 10, individual predictions and loss-influence ranges, are retained in Source code 1. The specifications are not independent replications.

##### Table S11A. First extinction to first retention

| Source priority | K | Raw baseline | Outcome | Raw RMSE | With config. RMSE | Persistence RMSE |
|---|---|---|---|---|---|---|
| Summary | 4 | F/P | Freezing | 31.092 | 41.694 | 29.684 |
| Summary | 4 | F/P | Inputs/min | 0.084 | 0.085 | 0.083 |
| Summary | 4 | Four-channel | Freezing | 30.253 | 41.629 | 29.684 |
| Summary | 4 | Four-channel | Inputs/min | 0.084 | 0.085 | 0.083 |
| Summary | 8 | F/P | Freezing | 31.092 | 45.348 | 29.684 |
| Summary | 8 | F/P | Inputs/min | 0.084 | 0.085 | 0.083 |
| Summary | 8 | Four-channel | Freezing | 30.253 | 45.606 | 29.684 |
| Summary | 8 | Four-channel | Inputs/min | 0.084 | 0.085 | 0.083 |
| Annotation | 4 | F/P | Freezing | 31.092 | 41.705 | 29.684 |
| Annotation | 4 | F/P | Inputs/min | 0.084 | 0.085 | 0.083 |
| Annotation | 4 | Four-channel | Freezing | 30.253 | 41.669 | 29.684 |
| Annotation | 4 | Four-channel | Inputs/min | 0.084 | 0.085 | 0.083 |
| Annotation | 8 | F/P | Freezing | 31.092 | 43.043 | 29.684 |
| Annotation | 8 | F/P | Inputs/min | 0.084 | 0.085 | 0.083 |
| Annotation | 8 | Four-channel | Freezing | 30.253 | 43.291 | 29.684 |
| Annotation | 8 | Four-channel | Inputs/min | 0.084 | 0.085 | 0.083 |

#### Table S11B. First extinction to renewal

| Source priority | K | Raw baseline | Outcome | Raw RMSE | With config. RMSE | Persistence RMSE |
|---|---|---|---|---|---|---|
| Summary | 4 | F/P | Freezing | 31.416 | 42.376 | 30.165 |
| Summary | 4 | F/P | Inputs/min | 0.589 | 0.596 | 0.583 |
| Summary | 4 | Four-channel | Freezing | 31.783 | 42.397 | 30.165 |
| Summary | 4 | Four-channel | Inputs/min | 0.590 | 0.596 | 0.583 |
| Summary | 8 | F/P | Freezing | 31.416 | 42.644 | 30.165 |
| Summary | 8 | F/P | Inputs/min | 0.589 | 0.597 | 0.583 |
| Summary | 8 | Four-channel | Freezing | 31.783 | 42.638 | 30.165 |
| Summary | 8 | Four-channel | Inputs/min | 0.590 | 0.598 | 0.583 |
| Annotation | 4 | F/P | Freezing | 31.416 | 42.878 | 30.165 |
| Annotation | 4 | F/P | Inputs/min | 0.589 | 0.595 | 0.583 |
| Annotation | 4 | Four-channel | Freezing | 31.783 | 42.913 | 30.165 |
| Annotation | 4 | Four-channel | Inputs/min | 0.590 | 0.595 | 0.583 |
| Annotation | 8 | F/P | Freezing | 31.416 | 45.286 | 30.165 |
| Annotation | 8 | F/P | Inputs/min | 0.589 | 0.593 | 0.583 |
| Annotation | 8 | Four-channel | Freezing | 31.783 | 45.445 | 30.165 |
| Annotation | 8 | Four-channel | Inputs/min | 0.590 | 0.593 | 0.583 |

#### Table S11C. Fourth extinction to second retention

| Source priority | K | Raw baseline | Outcome | Raw RMSE | With config. RMSE | Persistence RMSE |
|---|---|---|---|---|---|---|
| Summary | 4 | F/P | Freezing | 16.645 | 18.465 | 12.379 |
| Summary | 4 | F/P | Inputs/min | 1.668 | 2.239 | 2.257 |
| Summary | 4 | Four-channel | Freezing | 20.331 | 21.472 | 12.379 |
| Summary | 4 | Four-channel | Inputs/min | 1.827 | 2.292 | 2.257 |
| Summary | 8 | F/P | Freezing | 16.645 | 18.674 | 12.379 |
| Summary | 8 | F/P | Inputs/min | 1.668 | 2.166 | 2.257 |
| Summary | 8 | Four-channel | Freezing | 20.331 | 19.167 | 12.379 |
| Summary | 8 | Four-channel | Inputs/min | 1.827 | 2.221 | 2.257 |
| Annotation | 4 | F/P | Freezing | 16.645 | 14.662 | 12.379 |
| Annotation | 4 | F/P | Inputs/min | 1.668 | 2.532 | 2.257 |
| Annotation | 4 | Four-channel | Freezing | 21.540 | 17.913 | 12.379 |
| Annotation | 4 | Four-channel | Inputs/min | 1.872 | 2.407 | 2.257 |
| Annotation | 8 | F/P | Freezing | 16.645 | 23.616 | 12.379 |
| Annotation | 8 | F/P | Inputs/min | 1.668 | 2.232 | 2.257 |
| Annotation | 8 | Four-channel | Freezing | 21.540 | 24.048 | 12.379 |
| Annotation | 8 | Four-channel | Inputs/min | 1.872 | 2.328 | 2.257 |

#### Table S12. Added prediction of fecal-boli counts

Whole-acquisition comparisons use 98 acquisitions and task-window comparisons 148, each representing all 17 Study 2 animals. The outcome is always whole-acquisition boli count. Raw baselines include freezing, phase, group and cohort; four-channel also includes count rates. Configuration models add K-1 posterior fractions. Scores are held-out negative-binomial log scores averaged within animal, then equally across animals. Gain is with-configurations minus raw score; positive favors augmentation. Animals improved counts positive animal-level gains out of 17. Complete predictions, MAE, RMSE and descriptive influence ranges are in Source code 1. A difference in predictive score is not a test of biological equivalence.

##### Table S12A. Whole acquisition with duration offset

| Source priority | K | Raw baseline | Raw score | With config. score | Score gain | Animals improved |
|---|---|---|---|---|---|---|
| Summary | 4 | Freezing | -2.5127 | -2.5470 | -0.0343 | 9/17 |
| Summary | 4 | Four-channel | -2.5679 | -2.5903 | -0.0223 | 7/17 |
| Summary | 8 | Freezing | -2.5127 | -2.6034 | -0.0907 | 7/17 |
| Summary | 8 | Four-channel | -2.5679 | -2.6455 | -0.0776 | 7/17 |
| Annotation | 4 | Freezing | -2.5127 | -2.5455 | -0.0328 | 8/17 |
| Annotation | 4 | Four-channel | -2.5442 | -2.5748 | -0.0306 | 5/17 |
| Annotation | 8 | Freezing | -2.5127 | -2.6169 | -0.1042 | 8/17 |
| Annotation | 8 | Four-channel | -2.5442 | -2.6323 | -0.0881 | 4/17 |

##### Table S12B. Whole acquisition without duration offset

| Source priority | K | Raw baseline | Raw score | With config. score | Score gain | Animals improved |
|---|---|---|---|---|---|---|
| Summary | 4 | Freezing | -2.5094 | -2.5308 | -0.0214 | 9/17 |
| Summary | 4 | Four-channel | -2.5633 | -2.5724 | -0.0092 | 8/17 |
| Summary | 8 | Freezing | -2.5094 | -2.5948 | -0.0854 | 8/17 |
| Summary | 8 | Four-channel | -2.5633 | -2.6326 | -0.0693 | 7/17 |
| Annotation | 4 | Freezing | -2.5094 | -2.5298 | -0.0204 | 9/17 |
| Annotation | 4 | Four-channel | -2.5370 | -2.5532 | -0.0162 | 7/17 |
| Annotation | 8 | Freezing | -2.5094 | -2.5971 | -0.0877 | 9/17 |
| Annotation | 8 | Four-channel | -2.5370 | -2.6081 | -0.0711 | 4/17 |

#### Table S12C. Task window with duration offset

| Source priority | K | Raw baseline | Raw score | With config. score | Score gain | Animals improved |
|---|---|---|---|---|---|---|
| Summary | 4 | Freezing | -2.3162 | -2.3598 | -0.0436 | 5/17 |
| Summary | 4 | Four-channel | -2.3731 | -2.4422 | -0.0692 | 3/17 |
| Summary | 8 | Freezing | -2.3162 | -2.4298 | -0.1136 | 5/17 |
| Summary | 8 | Four-channel | -2.3731 | -2.5701 | -0.1970 | 3/17 |
| Annotation | 4 | Freezing | -2.3162 | -2.3642 | -0.0480 | 6/17 |
| Annotation | 4 | Four-channel | -2.3616 | -2.4223 | -0.0607 | 3/17 |
| Annotation | 8 | Freezing | -2.3162 | -2.4164 | -0.1003 | 3/17 |
| Annotation | 8 | Four-channel | -2.3616 | -2.5252 | -0.1636 | 2/17 |

#### Table S12D. Task window without duration offset

| Source priority | K | Raw baseline | Raw score | With config. score | Score gain | Animals improved |
|---|---|---|---|---|---|---|
| Summary | 4 | Freezing | -2.3255 | -2.3694 | -0.0439 | 5/17 |
| Summary | 4 | Four-channel | -2.3814 | -2.4490 | -0.0676 | 2/17 |
| Summary | 8 | Freezing | -2.3255 | -2.4467 | -0.1212 | 5/17 |
| Summary | 8 | Four-channel | -2.3814 | -2.5968 | -0.2154 | 2/17 |
| Annotation | 4 | Freezing | -2.3255 | -2.3732 | -0.0478 | 5/17 |
| Annotation | 4 | Four-channel | -2.3683 | -2.4273 | -0.0590 | 3/17 |
| Annotation | 8 | Freezing | -2.3255 | -2.4323 | -0.1068 | 3/17 |
| Annotation | 8 | Four-channel | -2.3683 | -2.5506 | -0.1823 | 2/17 |
