## Supplementary material for "Multimodal temporal modeling of behavioral reorganization across threat learning and extinction in rats": Source code 1: Additional outcome analyses, exact inputs and complete results: Figure_1.pdf

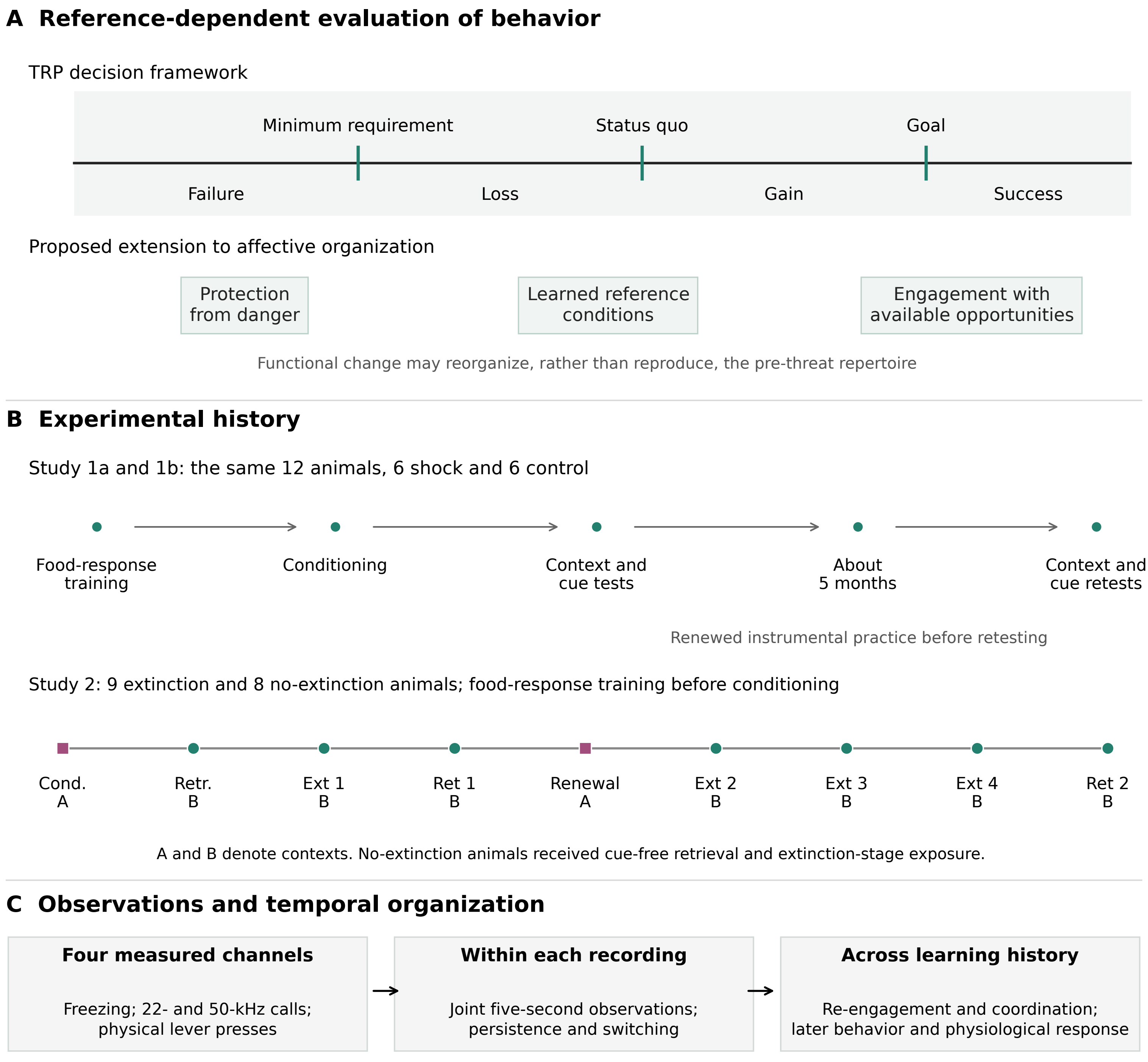

A Reference-dependent evaluation of behavior
TRP decision framework
Minimum requirement
Status quo
Goal
Failure
Loss
Gain
Success
Proposed extension to affective organization
Protection from danger
Learned reference conditions
Engagement with available opportunities
Functional change may reorganize, rather than reproduce, the pre-threat repertoire
B Experimental history
Study 1a and 1b: the same 12 animals, 6 shock and 6 control
Food-response training
Conditioning
Context and cue tests
About 5 months
Context and cue retests
Renewed instrumental practice before retesting
Study 2: 9 extinction and 8 no-extinction animals; food-response training before conditioning
Cond. A
Retr. B
Ext 1 B
Ret 1 B
Renewal A
Ext 2 B
Ext 3 B
Ext 4 B
Ret 2 B
A and B denote contexts. No-extinction animals received cue-free retrieval and extinction-stage exposure.
C Observations and temporal organization
Four measured channels
Freezing; 22- and 50-kHz calls; physical lever presses
Within each recording
Joint five-second observations; persistence and switching
Across learning history
Re-engagement and coordination; later behavior and physiological response
