## Supplementary figures and images for "Multimodal temporal modeling of behavioral reorganization across threat learning and extinction in rats"

### Figure_1.png

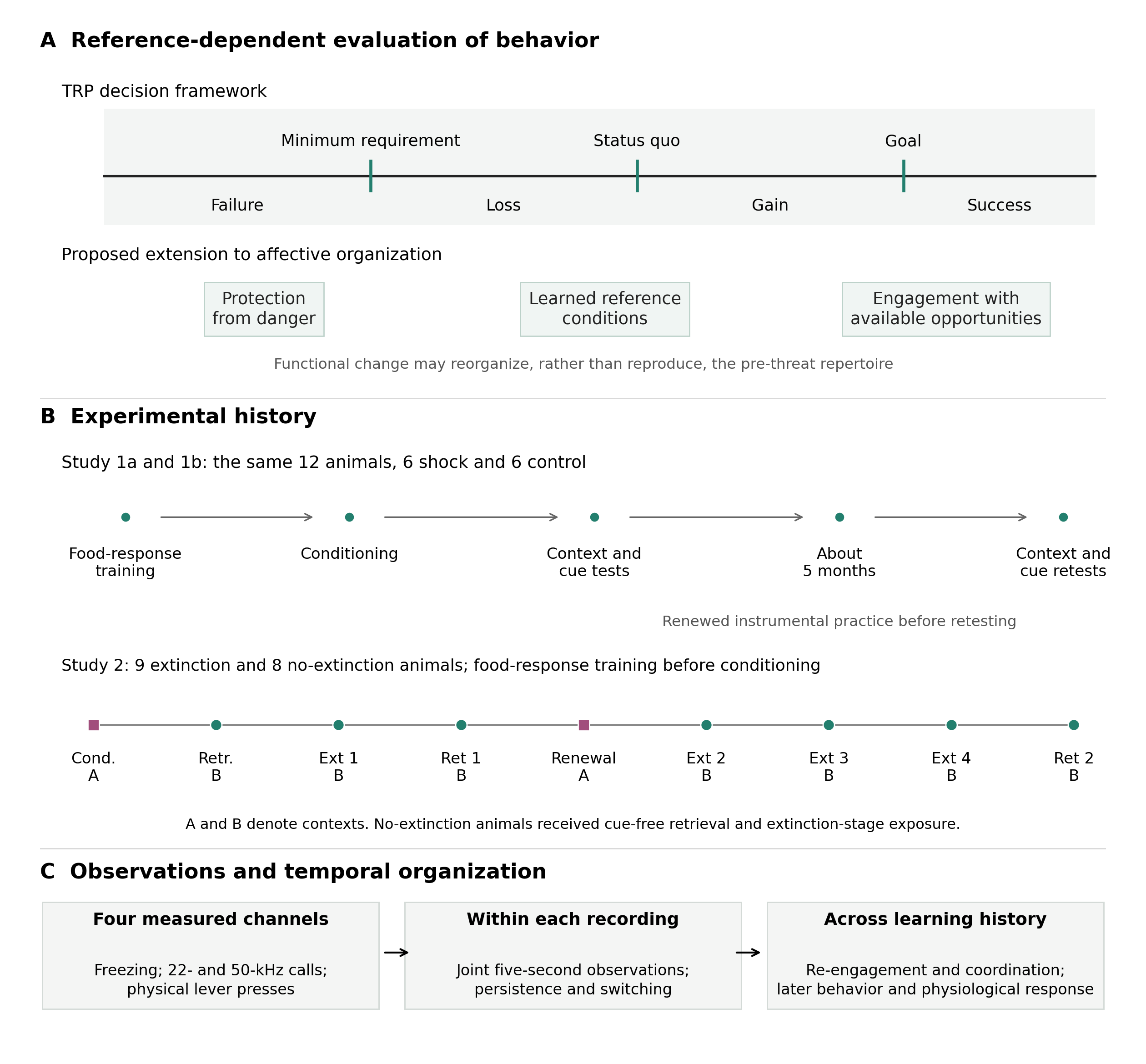
